# Rab11-FIP1C is dispensable for HIV-1 replication in primary CD4^+^ T cells but its role is cell-type-dependent in immortalized human T-cell lines

**DOI:** 10.1101/2022.06.03.494784

**Authors:** Melissa V. de Céspedes, Huxley K. Hoffman, Hannah Carter, Lacy M. Simons, Lwar Naing, Sherimay D. Ablan, Judd F. Hultquist, Schuyler B. van Engelenburg, Eric O. Freed

## Abstract

The HIV-1 envelope glycoprotein (Env) contains a long cytoplasmic tail harboring highly conserved motifs that direct Env trafficking and incorporation into virions and promote efficient virus spread. The cellular trafficking factor Rab11a family interacting protein 1C (FIP1C) has been implicated in the directed trafficking of Env to sites of viral assembly. In this study, we confirm that siRNA-mediated depletion of FIP1C in HeLa cells modestly reduces Env incorporation into virions. To determine whether FIP1C is required for Env incorporation and HIV-1 replication in physiologically relevant cells, CRISPR/Cas9 technology was used to knock out the expression of this protein in several human T-cell lines – Jurkat E6.1, SupT1 and H9 – and in primary human CD4^+^ T cells. *FIP1C* knock-out caused modest reductions in Env incorporation in SupT1 cells but did not inhibit virus replication in SupT1 or Jurkat E6.1 T-cell lines. In H9 cells, *FIP1C* knock-out caused a cell-density-dependent defect in virus replication. In primary CD4^+^ T cells, *FIP1C* knock-out had no effect on HIV-1 replication. Furthermore, HTLV-I transformed cell lines that are permissive for HIV-1 replication do not express FIP1C. Mutation of an aromatic motif in the Env cytoplasmic tail – Y_795_W – implicated in FIP1C-mediated Env incorporation impaired virus replication independently of FIP1C expression in SupT1, Jurkat E6.1, H9, and primary T cells. Together, these results indicate that while FIP1C may contribute to HIV-1 Env incorporation in some contexts, additional and potentially redundant host factors are likely required for Env incorporation and virus dissemination in T cells.

**Importance:** The incorporation of the HIV-1 envelope (Env) glycoproteins, gp120 and gp41, into virus particles is critical for virus infectivity. Gp41 contains a long cytoplasmic tail that has been proposed to interact with host cell factors, including the trafficking factor Rab11a family interacting protein 1C (FIP1C). To investigate the role of FIP1C in relevant cell types – human T-cell lines and primary CD4^+^ T cells – we used CRISPR/Cas9 to knock out FIP1C expression and examined the effect on HIV-1 Env incorporation and virus replication. We observed that in two of the T-cell lines examined (Jurkat E6.1 and SupT1) and in primary CD4^+^ T cells, FIP1C knockout did not disrupt HIV-1 replication, whereas FIP1C knockout reduced Env expression and delayed replication in H9 cells. The results indicate that while FIP1C may contribute to Env incorporation in some cell lines, it is not an essential factor for efficient HIV-1 replication in primary CD4+ T cells.

## Introduction

HIV-1 particle assembly takes place predominantly on the plasma membrane of infected host cells in a series of steps driven by the viral Gag polyprotein precursor. Although expression of Gag is sufficient for the formation of non-infectious virus-like particles (VLPs), the production of infectious virions requires expression and incorporation of the viral envelope glycoprotein (Env). HIV-1 Env is synthesized as a 160-kDa polyprotein precursor, gp160, on endoplasmic reticulum (ER)-associated ribosomes and is transported to the plasma membrane through the secretory pathway. During transport, gp160 trimerizes and is cleaved in the Golgi apparatus by host furin or furin-like enzymes to form two mature Env subunits, the surface glycoprotein subunit gp120 and the transmembrane subunit gp41.

The transmembrane Env subunits of most lentiviruses (including gp41 of HIV-1, HIV-2, and the related simian immunodeficiency viruses [SIVs]) have unusually long cytoplasmic tails (CTs) relative to those of other retroviruses, typically ∼150 amino acids in length [for reviews, see (1–3)]. These long CTs contain a variety of conserved motifs, including a Tyrosine-based, membrane-proximal YxxØ motif (where x is any residue and Ø is a hydrophobic residue), a di-Leucine motif at the C-terminus of the gp41 CT, and several aromatic motifs. Upon reaching the plasma membrane, Env is rapidly internalized via a clathrin-dependent pathway, in part mediated by direct interactions between the YxxØ and di-Leu motifs and the clathrin adaptor protein complexes AP-1 and AP-2 (4–12). Rapid Env internalization limits the amount of Env displayed on the cell surface, likely reducing the detection and elimination of infected cells by the host immune response (13). Env endocytosis also contributes to the low levels of Env trimers on HIV-1 particles compared to other retroviruses, with only approximately one dozen trimers per virion (14, 15). Although some of the internalized Env may be recycled back to the cell surface after endocytic uptake (16, 17), the role of Env recycling in Env incorporation is currently unclear.

The gp41 CT has also been reported to direct the sorting of Env in polarized epithelial cells and lymphocytes (18–21) and promotes Env accumulation at points of cell-cell contact referred to as the virological synapse (VS) (22). Transfer of virus across the VS is a highly efficient route of virus transmission, at least in cell culture, relative to cell-free particle infection (23–26). As a result of this highly efficient transmission, cell-cell transfer renders HIV-1 to be less sensitive to replication blocks and inhibitors (27–30), and mutations that enhance cell-cell transfer reduce the susceptibility of HIV-1 to antiretroviral drugs (31, 32).

Significantly, mutation of the YxxØ motif in the SIV_mac239_ gp41 CT allowed infected animals to control the viral infection, thereby reducing viral pathogenesis (33). Propagation of this virus *in vivo* led to the emergence of highly pathogenic revertants in which new endocytosis/trafficking motifs were acquired (34). Thus, the highly conserved endocytosis/trafficking motifs in lentiviral gp41 CTs play an important role in lentivirus propagation and pathogenesis.

In addition to playing a key role in Env trafficking and cell-surface expression, the long gp41 CT also mediates Env incorporation into virus particles [for reviews, see (1–3, 35)]. Truncation of the HIV-1 gp41 CT has little effect on Env incorporation in adherent cell lines such as HeLa, 293T, and in the MT-4 T-cell line, while the gp41 CT truncation mutant (36) is severely impaired for Env incorporation, particle infectivity, and virus spread in most T-cell lines, primary T cells, and monocyte-derived macrophages (37, 38). Mutation of the aforementioned highly conserved YxxØ and di-Leu motifs in the gp41 CT has also been reported to disrupt Env incorporation, particle infectivity, and virus propagation (33, 39, 40).

The MA domain of Gag plays a central role in HIV-1 Env incorporation [(36, 37, 41–44); for reviews, see (1, 3, 45)] and is thought to organize into hexamers of trimers on the inner leaflet of the viral envelope (46–48). Recent cryo-electron tomography data suggest that MA reorganizes during virus maturation, perhaps modulating Env function during the infection process (49). Recent structural studies have also provided insights into the conformation of the gp41 CT with respect to the membrane (50, 51), although how the gp41 CT might engage with the underlying MA lattice remains incompletely understood. It has been reported that the Gag lattice traps Env at the site of virus assembly, and that a MA mutation that blocks Env incorporation (36) prevents Env trapping in the assembling Gag lattice (52).

Rab proteins are members of the Ras superfamily of small GTPases [for reviews, see (53, 54)]. They function in a cascade-like fashion in the endolysosomal and secretory pathways to regulate membrane trafficking by promoting vesicle formation, movement, and fusion. Approximately 70 Rab proteins are expressed in human cells. Rab functions are mediated via the interaction of activated (GTP-bound) Rab proteins with a number of effectors that include tethering complexes, molecular motors, scaffolding proteins, and lipid kinases and phosphatases (55–57).

The cell-type-dependent role of the long gp41 CT in Env incorporation and virus replication supports the hypothesis that the CT interacts with host cell factors that regulate Env trafficking, surface expression, and virion incorporation. One such proposed gp41 CT-interacting host factor is Rab11-family interacting protein 1C (FIP1C; also known as Rab-coupling protein [RCP]). The FIPs are a family of effector proteins for Rab and ADP ribosylation factor (Arf) GTPases [for review see (58, 59)]. Depletion of FIP1C by using a small hairpin RNA (shRNA) was shown to reduce Env incorporation into HIV-1 particles in HeLa cells, in the H9 T-cell line, and in primary monocyte-derived macrophages (60, 61). In contrast, FIP1C did not affect the incorporation of the gp41 CT-truncated mutant CTdel144 (60–62). FIP1C depletion also delayed HIV-1 replication in H9 cells (60, 61). Depletion of Rab14, but not Rab11, also reduced Env incorporation in HeLa cells (60). Although a direct interaction between HIV-1 Env and FIP1C was not demonstrated, expression of full-length, but not CT-truncated, Env in HeLa cells induced a redistribution of FIP1C from a perinuclear compartment to the plasma membrane (60). The ability of HIV-1 Env to relocalize FIP1C was shown in a subsequent study to be dependent upon a Tyr-based motif – Y_795_W – in the gp41 CT, implicating this motif in FIP1C-dependent Env incorporation (61). Furthermore, overexpression of a dominant-negative fragment of FIP1C trapped WT but not CT-truncated or Y_795_W/SL-mutant Env in a perinuclear recycling compartment in HeLa cells (62). Together, these results supported the hypothesis that FIP1C-mediated recycling of Env is required for Env incorporation and implicated the Y_795_W motif in FIP1C-mediated Env recycling to virus assembly sites on the plasma membrane.

To evaluate the role of FIP1C in HIV-1 replication in T-cell lines and primary CD4^+^ T cells, we used CRISPR/Cas9 technology to knock-out (KO) the *FIP1C* gene. The results indicated variable and cell-type-dependent effects of *FIP1C* KO on HIV-1 replication in T-cell lines, and no defect in virus replication in primary CD4^+^ T cells. Mutation of the Y_795_W motif in the gp41 CT impaired Env incorporation irrespective of whether FIP1C was expressed. Overall, while the results support some role for FIP1C in Env incorporation, they indicate that other, or additional, host cell factors are likely involved. The results also indicate that the Y_795_W motif plays a role in Env incorporation and virus propagation that is independent of FIP1C.

## Results

### Knockdown of FIP1C expression in HeLa cells modestly reduces virion incorporation of WT Env but not incorporation of CTdel144 or Y_795_W/SL Env mutants

HIV-1 replication in most T-cell lines is abrogated by truncation of the gp41 CT (37, 38, 63, 64). Several studies have shown that mutating a hydrophobic motif, Tyr_795_Trp (Y_795_W), located in the LLP3 domain of the gp41 CT, to Ser-Leu (Y_795_W/SL; Figure. 1A), significantly reduces Env incorporation (39, 61, 62). This motif has been implicated in FIP1C-mediated trafficking of Env to viral assembly sites (60–62).

**Figure 1.**
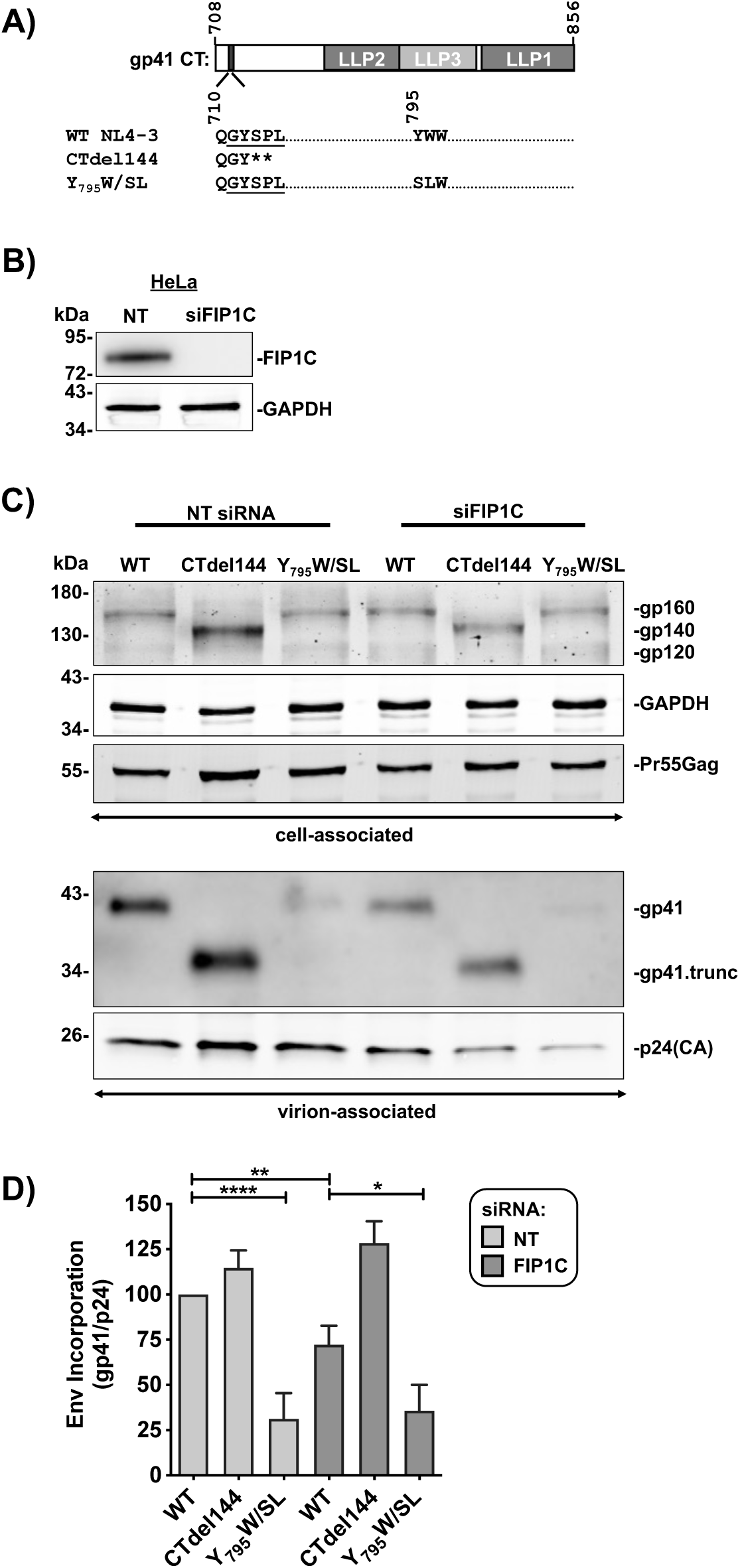
Knockdown of FIP1C expression in HeLa cells modestly reduces virion incorporation of WT Env but not incorporation of CTdel144 or Y_795_W/SL Env mutants. (**A**) Schematic representation of the NL4-3 gp41 CT and gp41 CT genotypes used in this study. The lentiviral lytic peptide (LLP) domains are indicated in gray boxes, and the highly conserved tyrosine endocytosis motif is indicated with a shaded black rectangle. The numbers above the gp41 CT schematic indicate the first and last amino acid positions of the gp41 CT. The number below the gp41 CT indicates the position of the QGYSPL sequence and the YW^795^ motif. The CTdel144 mutant was generated by introducing two stop codons in the highly conserved tyrosine endocytosis motif, GYSPL, resulting in a CT of 4 amino acids (36). (**B**) Cell lysates HeLa cells were treated with either NT or FIP1C-targeting siRNA were analyzed by Western blotting for FIP1C expression. (**C**) HeLa cells treated with siRNA were transfected with either WT or CTdel144 proviral DNA and samples were collected 48 h post-transfection. Western blotting was performed on the virus fraction to detect gp41 and p24(CA) using equal amounts of WT and CTdel144 viral lysates. CTdel144 was indicated by the shift in the molecular weight of Env and labelled as gp41.trunc and gp160.trunc. (**D**) Env incorporation, calculated by determining the ratio of virion-associated gp41 to p24(CA) relative to the WT condition, is represented. The bar graphs show the mean values ± standard deviation from three independent experiments. Statistical significance was assessed by one-way ANOVA and Tukey’s multiple-comparison test.

To begin our investigation into the role of FIP1C in the incorporation of WT and mutant Env, HeLa cells were treated with siRNA targeting FIP1C (siFIP1C) or a nontargeting control (NT; Figure 1B) and were transfected with the pNL4-3 molecular clone encoding WT, CT-truncated (CTdel144) (36), or Y_795_W/SL Env (Figure 1C). The siRNA-mediated knockdown was highly efficient, as no FIP1C was detectable in siFIP1C-treated cells (Figure 1B). Cell and virus fractions were analyzed by western blotting to detect virally encoded proteins (Figure 1C). FIP1C knockdown (KD) resulted in an approximately 25% reduction in WT Env incorporation without any reduction in the incorporation of CTdel144 or Y_795_W/SL Env (Figure 1D). While the effects of FIP1C KD on WT Env incorporation were modest, this result is consistent with the previously described role for FIP1C in HIV-1 Env incorporation in HeLa cells (60–62).

### FIP1C is expressed in non-HTLV transformed T-cell lines

FIP1C expression in human T-cell lines has been reported in H9 (60) and CEM-SS T-cell lines (65). To measure FIP1C protein expression in cell lines commonly used in the study of HIV-1 replication, we performed western blotting using a range of cell lines (Figure 2A). We also examined the expression levels of Rab14, which has been implicated in Env incorporation (60). HeLa cells were used as a positive control for FIP1C expression. FIP1C expression in HeLa and 293T was approximately equivalent, while FIP1C protein levels in the T-cell lines tested – SupT1, C8166, and M8166 – varied. Consistent with a previous report (65), and despite being highly permissive for HIV-1 replication (64, 66), MT-4 cells did not express detectable levels of FIP1C. All four T-cell lines expressed Rab14. To further interrogate the defect in FIP1C gene expression in MT-4 cells, RNA-seq was performed to measure RNA levels of the FIP1C isoform (Figure 2B). RNA levels closely mirrored protein levels (Figure 2A and B), indicating that the defect in FIP1C protein expression in MT-4 cells is due to a lack of FIP1C RNA expression.

**Figure 2.**
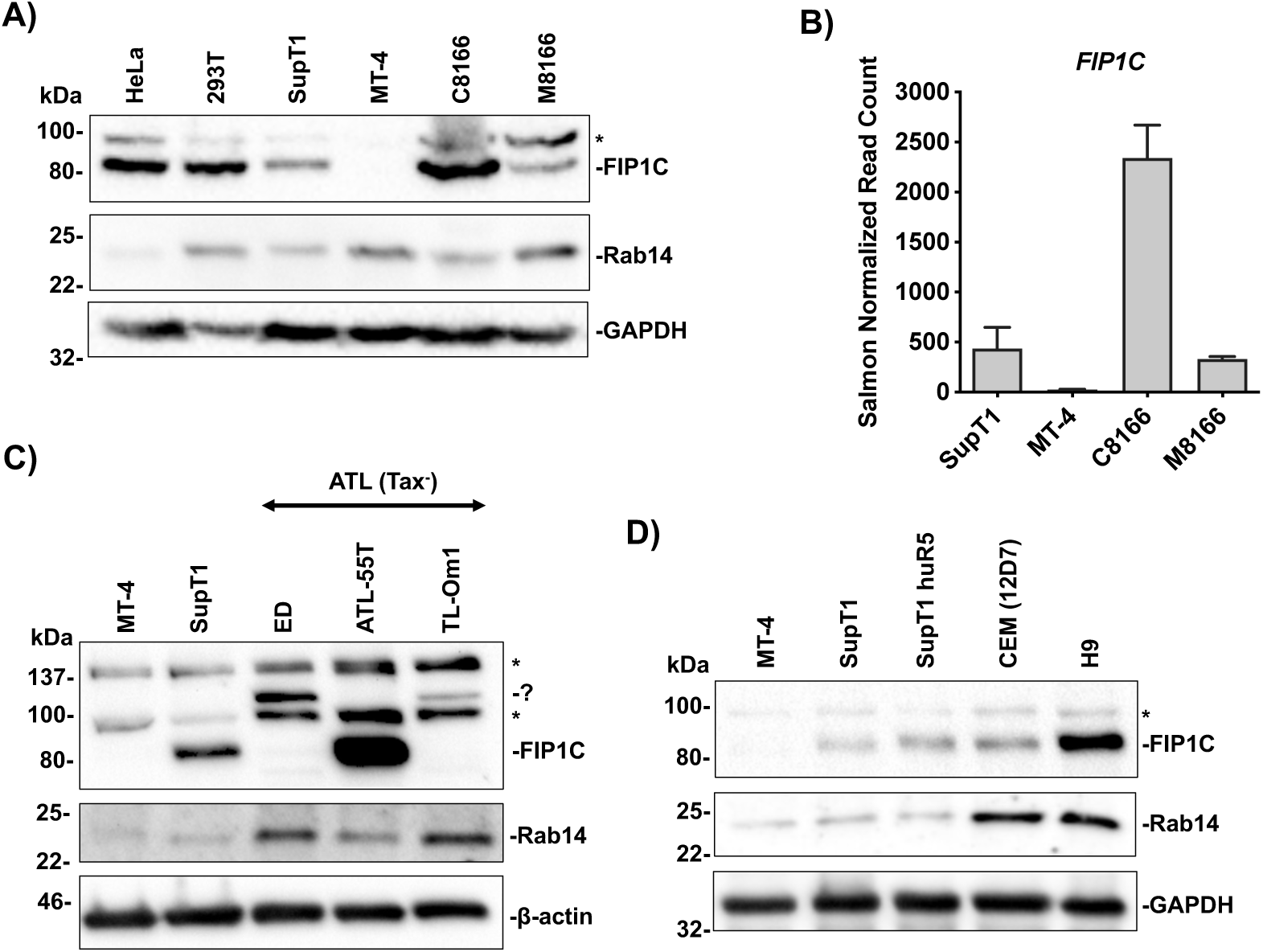
FIP1C is expressed in non-HTLV transformed T-cell lines. (**A, C–D**) Equivalent amounts of cells were used to generate the cell lysates and equivalent volumes of lysate were analyzed by Western blotting for FIP1C and Rab14 and a loading control, either GAPDH or β-actin. Proteins were detected via chemiluminescence. (**B**) RNA levels were determined by Illumina RNA-seq and RPKM values for each cell line were graphed. Western blot analysis is representative of three independent experiments. RNA-seq values are representative of duplicate RNA-seq analysis. A background band is denoted by a “*” and potential FIP isoforms, or background bands, are indicated with a “?”

MT-4 is an HTLV-I transformed cell line (67) that expresses HTLV-I Tax (64, 66, 68). To examine whether FIP1C and Rab14 expression may be influenced by HTLV-I transformation and determine their expression levels across previously un-evaluated T-cell lines permissive to HIV-1 replication, FIP1C and Rab14 expression in other HTLV-1-transformed T-cell lines was evaluated (Figure 2C). Western blotting was performed to measure FIP1C and Rab14 expression in several adult T-cell leukemia/lymphoma (ATL) cells that express little or no detectable Tax protein (Tax^−^) (64), though some of these cell lines (e.g., TL-Om1 and ED) were permissive to HIV-1 replication (69). The FIP1C-deficient MT-4 and FIP1C-expressing SupT1 were included as negative and positive controls, respectively. Two of the ATL Tax^−^ cell lines, ED and TL-Om1, did not express FIP1C but instead expressed an additional band of larger molecular weight of approximately 95 kDa. This additional band, which could be a background band or an additional FIP isoform, was not present in the MT-4 or SupT1 samples. ATL-55T cells expressed high levels of FIP1C and did not express this additional band of larger molecular weight. FIP1C expression was not linked to Rab14 protein expression, as all tested ATL Tax^−^ cell lines expressed Rab14, albeit at varying levels (Fig. 2C).

To assess FIP1C and Rab14 expression across non-ATL T-cell lines, western blotting was performed on SupT1huR5, CEM(12D7), and H9 cell lysates (Figure 2D). FIP1C and Rab14 expression levels varied amongst these cell lines, with FIP1C and Rab14 expression highest in H9. Altogether, these data indicate that FIP1C is expressed in all non-HTLV-I transformed human T-cell lines tested, while FIP1C expression is deficient in 3 out of the 6 HTLV-I-transformed cell lines tested. Additionally, Rab14 was expressed in all T-cell lines tested. Importantly, despite being highly permissive for HIV-1 replication, the MT-4 T-cell line does not express detectable levels of FIP1C. Altogether, these data show that FIP1C is not required for T-cell permissivity to HIV-1 replication.

### FIP1C KO in Jurkat E6.1 cells does not impair Env incorporation or viral replication kinetics

A role for FIP1C in HIV-1 replication in the H9 T-cell line was demonstrated by using shRNA-mediated knock-down of FIP1C expression (60). To determine whether FIP1C is required for Env incorporation in the Jurkat E6.1 T-cell line, CRISPR/Cas9 gene editing was used to generate two single-cell-derived FIP1C knock-out (KO) clones, denoted KO #6 and #9. No FIP1C protein expression was detected in either KO clone by western blotting (Figure 3A). Env incorporation in the FIP1C KO cell lines was compared to that in the parental cell line (Figure 3A). FIP1C KO did not alter Env incorporation in KO #6, but the KO #9 clone incorporated 200% more WT Env than parental cells or KO clone #6 (Figure 3A-B). Clone #9 also displayed an overall increase in viral protein expression (Gag and Env). The Y_795_W/SL mutant displayed reduced Env incorporation relative to WT, consistent with a role for this motif in Env incorporation (39, 61, 62). However, as in the case of WT Env, FIP1C KO did not reduce incorporation of this Env mutant (Fig. 3A).

**Figure 3.**
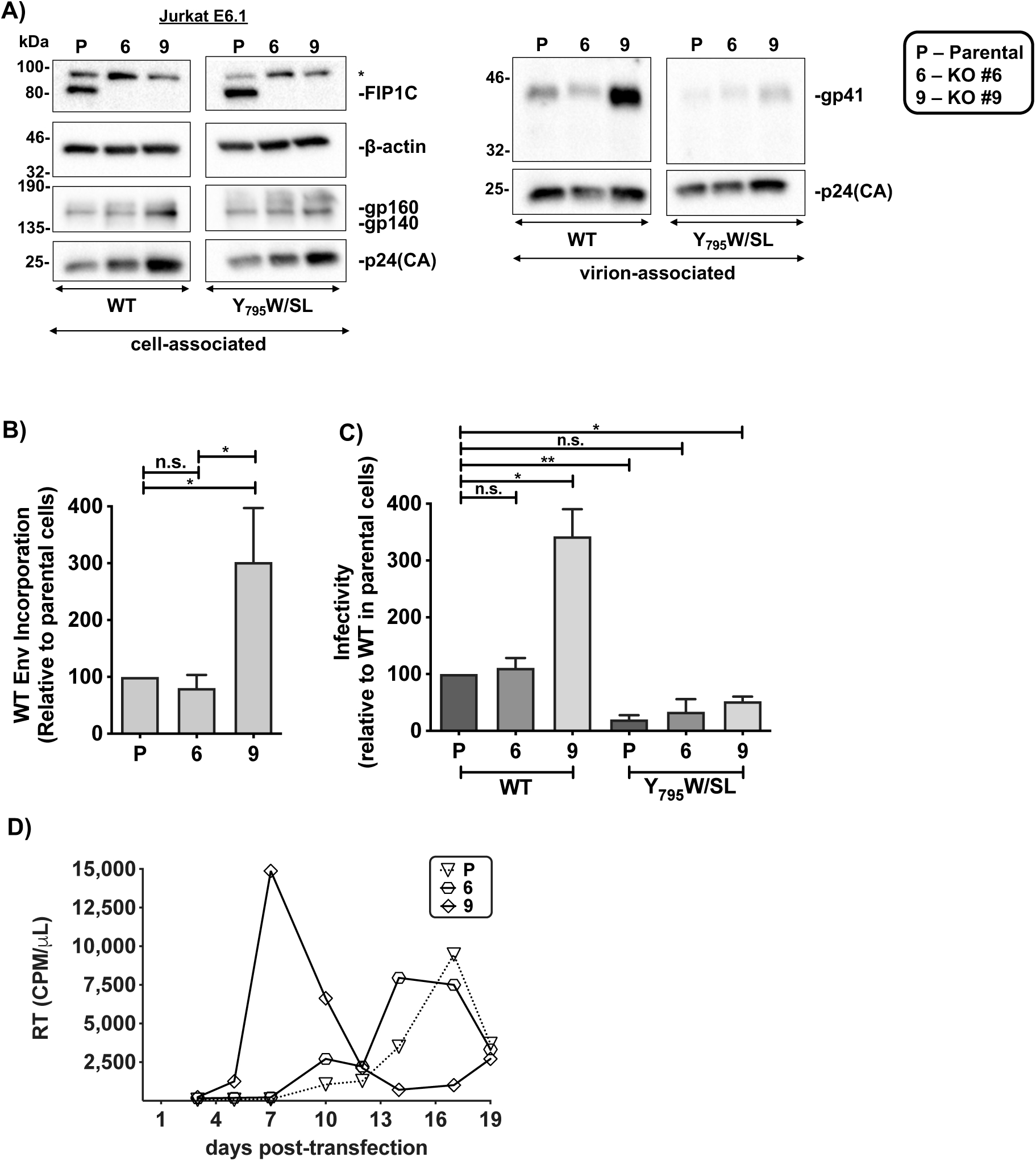
FIP1C KO in Jurkat E6.1 cells does not impair Env incorporation or viral replication kinetics. (**A**) Cells were transduced with RT-normalized VSV-G pseudotyped NL4-3 encoding either WT, CTdel144, or YW_795_SL Env. Western blotting was performed on the cell fraction to detect FIP1C, Gag precursor Pr55Gag, p24(CA), and gp160 and on the virus fraction to detect gp41 and p24(CA) using equal amounts of viral lysates. gp41.trunc indicates the position of the truncated gp41, CTdel144. A background band is denoted by a “*” (**B**) Relative Env incorporation was calculated by determining the ratio of virion-associated gp41 to p24(CA) relative to WT. (**C**) Viral supernatants were RT normalized used to infect TZM-bl cells. Luciferase values were then used to determine the relative infectivity of virus produced from each Jurkat E6.1 T-cell line. The graph shows the mean ± standard deviation of virions produced from KO cell lines for WT and YW_795_SL Env genotypes. (**D**) Replication curves for spreading infection of WT NL4-3 in parental and KO Jurkat E6.1 cell lines are shown. Cell lines were transfected with the proviral clone and split 1/3 and replenished with fresh RPMI-10% FBS every 2–3 days. An aliquot of the cell culture supernatant was reserved for analysis of HIV-1 RT activity at each time point. Replication kinetics are representative of 3 independent experiments. Statistical significance was assessed by Student’s t-test. n.s. indicates no statistical significance.

To determine whether FIP1C KO affects viral infectivity independently of virion Env content, cell-free infectivity of released virus was determined (Figure 3C). Virus particles produced from Jurkat E6.1 KO clone #6 displayed WT levels of infectivity, whereas, consistent with increased Env content, virus produced from KO clone #9 were 250% more infectious than WT virions produced from parental cells. The infectivity of Y_795_W/SL mutant virions produced from the KO clones was either not different from the infectivity of this mutant produced in parental cells (clone #6) or displayed increased infectivity (clone #9, consistent with the increase in Env incorporation in this clone). Thus, while the Y_795_W/SL mutant displayed reduced infectivity relative to WT in the Jurkat E6.1 cell line, this reduced infectivity was not linked to FIP1C expression.

To compare the replication of WT HIV-1 in the parental and KO Jurkat E6.1 clones, cells were transfected with WT pNL4-3 and replication kinetics were monitored by reverse transcriptase (RT) assay (Figure 3D). WT HIV-1 replication in KO clone #6 peaked three days earlier compared to parental Jurkat E6.1 cells. In KO clone #9, virus replicated with substantially faster kinetics than in parental cells, consistent with increased viral gene expression, higher Env content, and increased infectivity of virus produced in this cell clone. Altogether, these findings demonstrate that FIP1C is not required for Env incorporation or HIV-1 replication in the Jurkat E6.1 T-cell line. The increases in viral gene expression relative to parental cells in KO #9 also highlight the potential pitfalls associated with single-cell-derived clones (see Discussion).

### FIP1C KO in the SupT1 T-cell line modestly reduces viral protein expression and Env incorporation but does not affect virus replication kinetics

The data presented in Fig. 3 indicate that FIP1C is not required for HIV-1 Env incorporation or virus replication in the Jurkat E6.1 T-cell line. To examine the role of FIP1C in another T-cell line, we generated FIP1C KOs in the CCR5-expressing SupT1 T-cell line SupT1huR5. To avoid the clonal variability observed with the single-cell-derived Jurkat E6.1 KO clones (see above), we generated a polyclonal FIP1C KO SupT1huR5 cell line by. A cell line expressing Cas9, but not the gRNA, was also generated for use as a control (CTRL). FIP1C expression was undetectable by western blotting in the FIP1C KO SupT1huR5 cell line but readily detectable in the Cas9-expressing CTRL line (Figure 4A).

**Figure 4.**
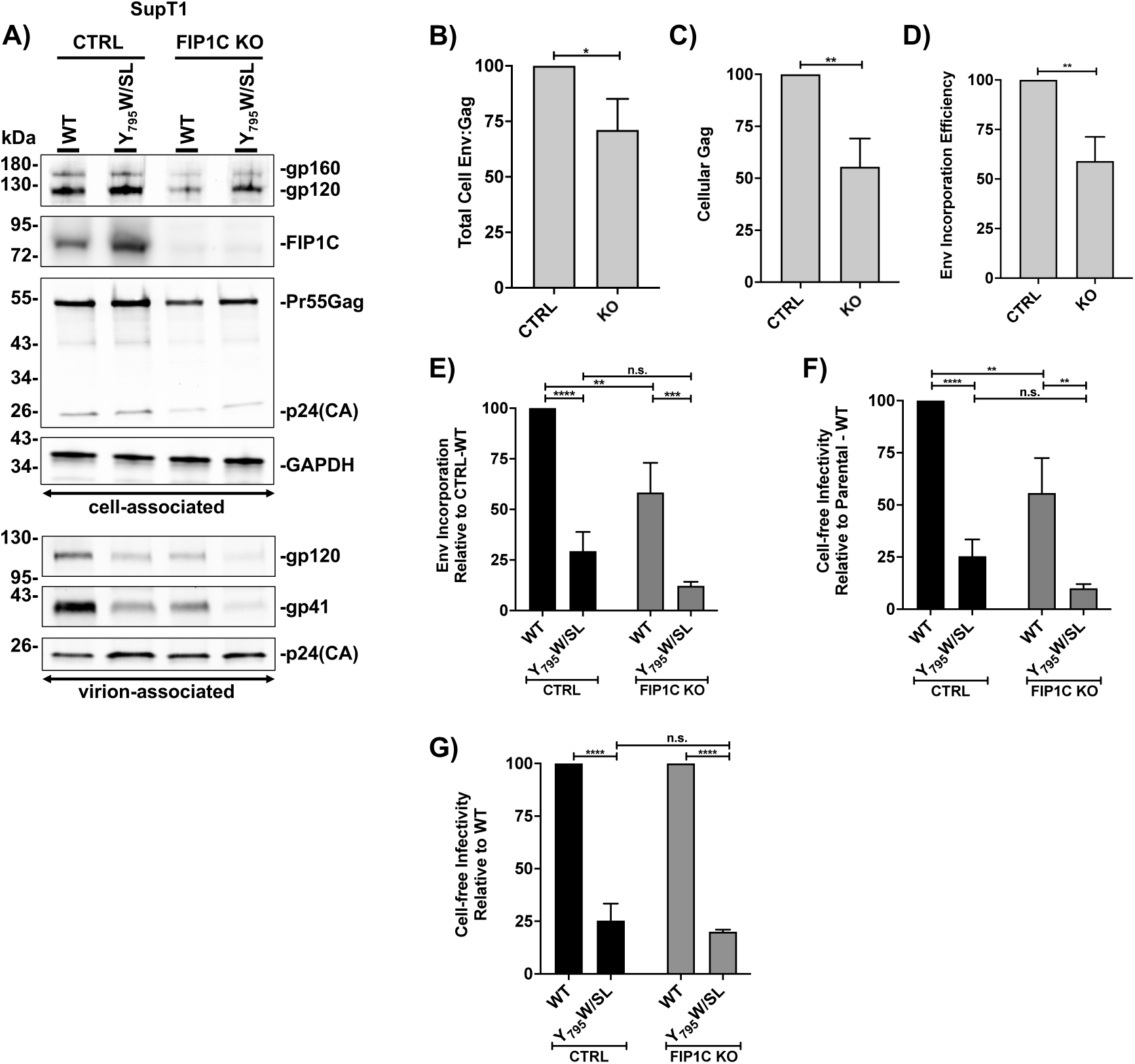
FIP1C KO in the SupT1 T-cell line modestly reduces viral protein expression and Envincorporation. (**A**) Cells were transduced with RT-normalized VSV-G pseudotyped NL4-3 encoding either WT or YW_795_SL Env. Western blot analysis was performed on the cell fraction to detect FIP1C and viral products and on the virus fraction to detect viral products using equal amounts of viral lysates. Various endpoints were measured from the band intensities on the Western blots: (**B**) Cellular Env:Gag Ratio = (gp120_c_+ gp160_c_)/[Pr55(Gag)_c_+p24(CA)_c_], (**C**) Cellular Gag Expression = [Pr55(Gag)_c_+p24(CA)_c_]/GAPDH_c_ (**D**) Env incorporation efficiency = [gp41_v_/p24(CA)_v_]/[gp120_c_+ gp160_c_], (**E**) Env incorporation = gp41_v_/p24(CA)_v_, where “c” is cell and “v” is virus. (**F–G**) Viral supernatant were RT-normalized and a serial dilution of virus was used to infect TZM-bl cells. Luciferase values were then used to determine the relative infectivity of virus produced from each T-cell line. The bar graphs show the mean values of virions produced from the CTRL and the KO cell lines for WT and YW_795_SL Env genotypes from both cell lines relative to (**F**) WT virions produced by CTRL cells or (**G**) WT produced by either cell line. Data are representative of the mean of three independent experiments ± standard deviation. Statistical significance was assessed by Student’s t-test. n.s. indicates no statistical significance.

To determine the effect of FIP1C KO on Env incorporation, CTRL and FIP1C KO SupT1huR5 cells were transduced with VSV-G-pseudotyped NL4-3 encoding either WT or Y_795_W/SL Env. Cell and viral lysates were analyzed for Gag and Env expression by western blotting (Figure 4A). The cellular Env:Gag ratio (Figure 4B) and the cellular Gag expression relative to GAPDH of WT virus in FIP1C KO cells relative to CTRL cells were determined by western blot (Figure 4C) . The results indicated that the ratio of Env:Gag and Gag expression overall were lower in FIP1C KO cells compared to CTRL cells. Reductions in Env incorporation (the ratio of gp41_v_/p24[CA]_v_, where v denotes virion-associated protein) were likewise observed with reduced Env incorporation partially due to reduced Env expression in the FIP1C KO cells. There was an approximately 33% reduction in WT Env incorporation efficiency, defined as (gp41_v_/p24[CA]_v_)/(gp120_c_+gp160_c_), where c and v denote cell-and virion-associated protein, respectively (Figure 4D).

We also compared the Env incorporation efficiency for the Y_795_W/SL mutant in SupT1 FIP1C KO vs CTRL cells. The Y_795_W/SL mutant exhibited a significant, and comparable, reduction in Env incorporation in both FIP1C KO and CTRL cells (Figure 4E). Cell-free, single-cycle infectivity of the virions generated in these cells was also calculated. TZM-bl cells were infected with RT-normalized virus and luciferase expression was quantified and normalized to CTRL-WT conditions (Figure 4F) and relative to WT conditions for each cell line (Figure 4G). The cell-free infectivity data paralleled the virion Env incorporation levels (Figure 4F compared to 4E). When values were normalized to WT conditions for each cell line, the defect imposed by the Y_795_W/SL mutant was essentially identical in CTRL and FIP1C KO cell lines (Figure 4G). Altogether, these data indicate that mutation of the Y_795_W motif caused a significant reduction in Env incorporation that is independent of FIP1C expression and that FIP1C KO caused modest effects on Env expression that contributed to the observed reductions in Env incorporation.

We next examined the effect of FIP1C KO on NL4-3 replication kinetics in SupT1 cells. The CTRL and FIP1C KO cells were transfected with WT, a mutant not expressing Env [Env(-)], or Env Y_795_W/SL NL4-3 proviral clones and virus replication was monitored by RT assay (Figure 5A). As expected, the Env(-) NL4-3 derivative did not replicate in either CTRL or KO cells. The NL4-3 WT clone replicated in CTRL and FIP1C KO cells with essentially identical kinetics, indicating that FIP1C is not required for the spread of NL4-3 in SupT1huR5 cultures. The Y_795_W/SL mutant replicated with similarly delayed kinetics in CTRL and KO cells, indicating that the replication defect exhibited by the Y_795_W/SL mutant in this T-cell line is unrelated to FIP1C expression.

**Figure 5.**
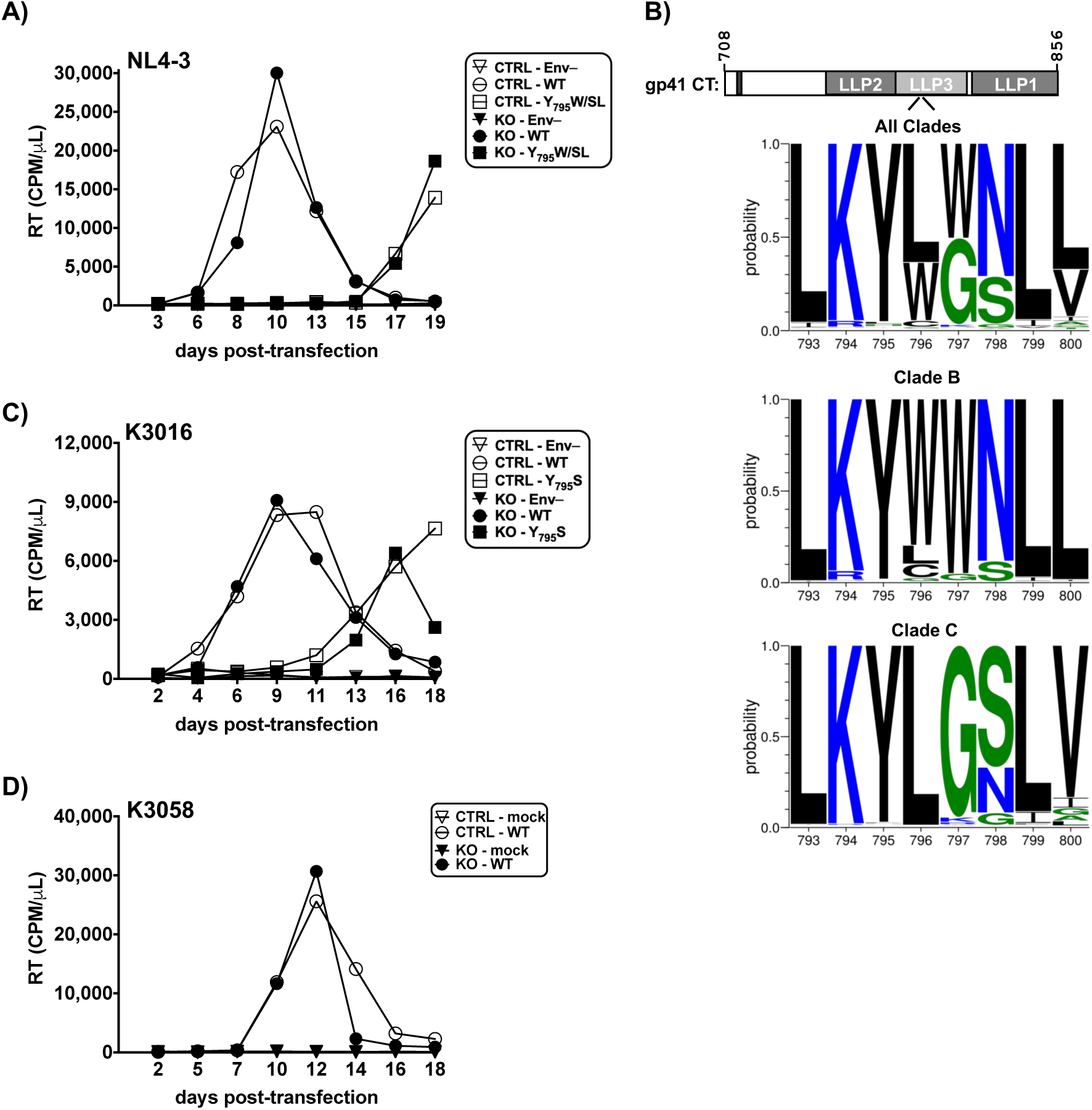
FIP1C KO in SupT1 does not affect viral replication kinetics of NL4-3 and clade C transmitted/founder HIV-1 strains K3016 and K3058. (**A, C–D**) Replication curves for spreading infection of WT and YW_795_SL NL4-3, WT and Y_795_S K3016, and WT K3058 are shown. NL4-3 replications were performed in SupT1 and K3016 and K3058 replications were performed in SupT1huR5. Cell lines were transfected with the proviral clone and split 1/3 and replenished with fresh RPMI-10% FBS every 2–3 days. An aliquot of the cell culture supernatant was reserved for analysis of HIV-1 RT activity at each time point. Data are representative of three independent experiments. (**B**) Schematic representation of the NL4-3 gp41 CT and gp41 CT genotypes used in this study. The lentiviral lytic peptide (LLP) domains are indicated in gray boxes, and the highly conserved tyrosine endocytosis motif is indicated with a shaded pink rectangle. The numbers above the gp41 CT schematic indicate the first and last amino acid positions of the gp41 CT. The frequency of amino acids by position of HIV-1 sequences in the Los Alamos database are shown for All clades, Clade B, or Clade C.

The Y_795_W motif is highly conserved amongst subtype B viruses, whereas only Y_795_ is conserved in subtype C viruses, with position 796 frequently a Leu rather than a Trp (Figure 5B). Among the >6,000 subtype B Env amino acid sequences in the Los Alamos National Laboratory (LANL) database at the time of this analysis (70) Y_795_ is 98.24% conserved, and amongst sequences across all subtypes Y_795_ is 96.74% conserved. Due to the high degree of conservation of Y_795_ and absence of W at position 796 in subtype C viruses, a Y_795_S mutant was generated in the subtype C transmitted/founder strain K3016. CTRL and FIP1C KO SupT1huR5 cells were transfected with K3016 proviral clones expressing either WT Env or the Y_795_S mutant and virus replication was monitored (Figure 5C). The WT K3016 clone replicated with identical kinetics in CTRL and FIP1C KO cultures, indicating that FIP1C is not required for the spread of K3016 in SupT1huR5 cultures. The Y_795_S mutant exhibited similarly delayed replication in both the CTRL and FIP1C KO cells, indicating that the replication defect imposed by the Y_795_S mutation in the context of the K3016 isolate is unrelated to FIP1C expression. The effect of FIP1C KO on the replication of another subtype C transmitted/founder virus, K3058, was also evaluated (Figure 5D). FIP1C KO did not alter virus replication kinetics for this isolate. Altogether, these data indicate that FIP1C expression is not required for the spread of NL4-3, K3016, or K3058 strains in the SupT1huR5 T-cell line. Furthermore, mutation of the Y_795_W motif in NL4-3 and Y_795_ in K3016 resulted in replication defects that were unaffected by FIP1C KO, similar to NL4-3 and the NL4-3 Y_795_W/SL Env mutant.

### FIP1C KO in H9 cells results in reduced viral protein expression and Env incorporation, and attenuated virus replication

As mentioned above, shRNA-mediated knock-down of FIP1C in H9 cells was demonstrated to delay HIV-1 replication (60). To corroborate these results by using CRISPR/Cas9 KO technology, a polyclonal FIP1C KO H9 cell line was generated by using the same method employed for SupT1huR5. In this case, we used the parental cells (Parental) as a control. FIP1C was detected by both fluorescence- and chemiluminescence-based western blotting; the latter being the more sensitive detection method. FIP1C was not detected by either method (Figure 6A), indicating a highly efficient KO. Due to the utilization of different blocking buffers using these two detection methods, the pattern of background bands was different (as indicated by the asterisks).

**Figure 6.**
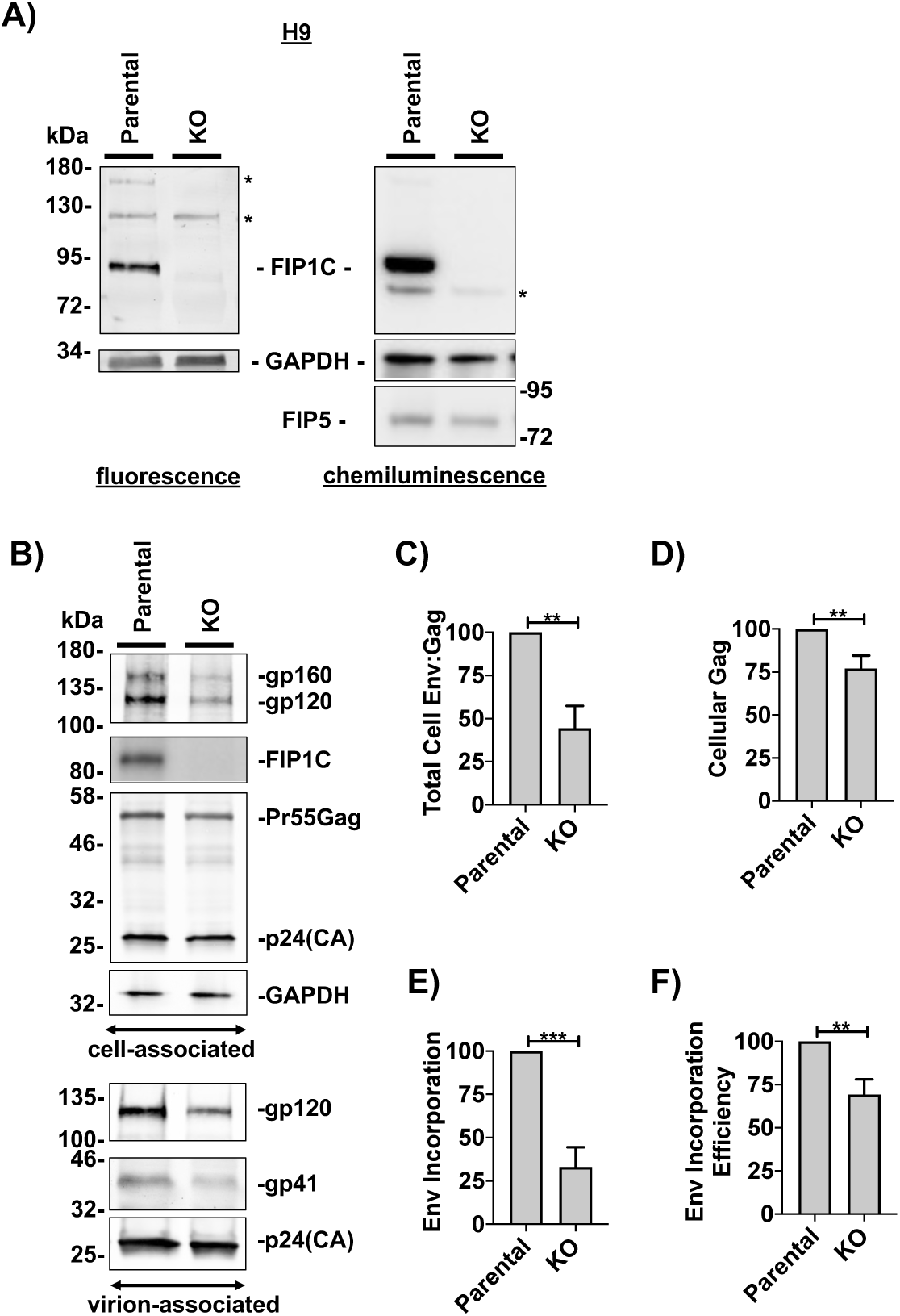
FIP1C KO in H9 cells results in reduced viral protein expression and Env incorporation. (**A**) An equivalent number of cells were lysed to generate cell lysates for analysis and an equivalent volume of cell lysates were analyzed by Western blot analysis. Samples were analyzed for FIP1C expression by fluorescence and chemiluminescence (see Materials and Methods). A background band is denoted by a “*”. (**B**) Cells were transduced with RT-normalized VSV-G pseudotyped NL4-3 encoding WT Env. Western blot analysis was performed on the cell fraction to detect FIP1C and viral products and on the virus fraction to detect viral products using equal amounts of viral lysates. Various endpoints were measured from the band intensities on the Western blots: (**C**) Cellular Env:Gag Ratio = (gp120_c_+ gp160_c_)/[Pr55(Gag)_c_+p24(CA)_c_], (**D**) Cellular Gag Expression = [Pr55(Gag)_c_+p24(CA)_c_]/GAPDH_c_ (**E**) Env incorporation efficiency = [gp41_v_/p24(CA)_v_]/[gp120_c_+ gp160_c_], (**F**) Env incorporation = gp41_v_/p24(CA)_v_, where “c” is cell and “v” is virus. Data are representative of the mean of three independent experiments ± standard deviation. Statistical significance was assessed by student T-test.

We next assessed the effect of FIP1C KO on Env incorporation in H9 cells. Parental and FIP1C KO H9 cells were transduced with VSV-G-pseudotyped NL4-3 encoding WT Env. Cell and viral fractions were analyzed by western blotting (Figure 6B). From these blots, the cellular Env:Gag ratio (Figure 6C) and cellular Gag expression relative to GAPDH (Figure 6D) in FIP1C KO H9 cells relative to Parental cells were determined. We found that the Env:Gag ratio, Env expression, and Gag expression were lower in FIP1C KO cells compared to Parental cells, similar to the phenotype observed in SupT1huR5 KO cells. The effect on Env expression was greater than the effect on Gag expression. To quantify Env incorporation in the FIP1C KO cells, the percent Env incorporation relative to the Parental cells was calculated as was done with SupT1 cells. FIP1C KO resulted in an approximately 66% reduction in Env content in virions (Figure 6E). To determine whether the reduced Env:Gag ratio contributed to the reduced virion Env content, we calculated the Env incorporation efficiency in FIP1C KO versus Parental cells (Figure 6F) as was done with SupT1 cells. After performing this normalization for cell-associated Env levels, we calculated a small (∼30%) but statistically significant reduction in Env incorporation efficiency, which indicates that the Env incorporation defect in the H9 FIP1C KO cell line is at least partially due to reduced Env expression.

To determine the effect of FIP1C KO on HIV-1 replication in H9 cells, Parental and KO cells were transfected with NL4-3 proviral clones encoding either Env(-), WT, or Y_795_W/SL Env. Virus replication was monitored by supernatant RT levels (Figure 7A-B). In both the Parental and FIP1C KO cells, Env(-) NL4-3 did not replicate, as anticipated. The NL4-3 WT clone peaked on day 9 in the Parental line but exhibited attenuated replication in the FIP1C KO cells. The Y_795_W/SL mutant exhibited a dampened and delayed replication curve in the Parental line and was highly defective for spreading infection in the FIP1C KO line. Therefore, in contrast to the Jurkat E6.1 and SupT1huR5 cell lines, FIP1C KO resulted in replication defects in the H9 cell line. We consistently observed less efficient replication of WT NL4-3 in H9 FIP1C KO cells compared to parental H9 and the other T-cell lines tested, as determined by the lower levels of RT activity at the peak of replication in H9 FIP1C KO cells.

**Figure 7.**
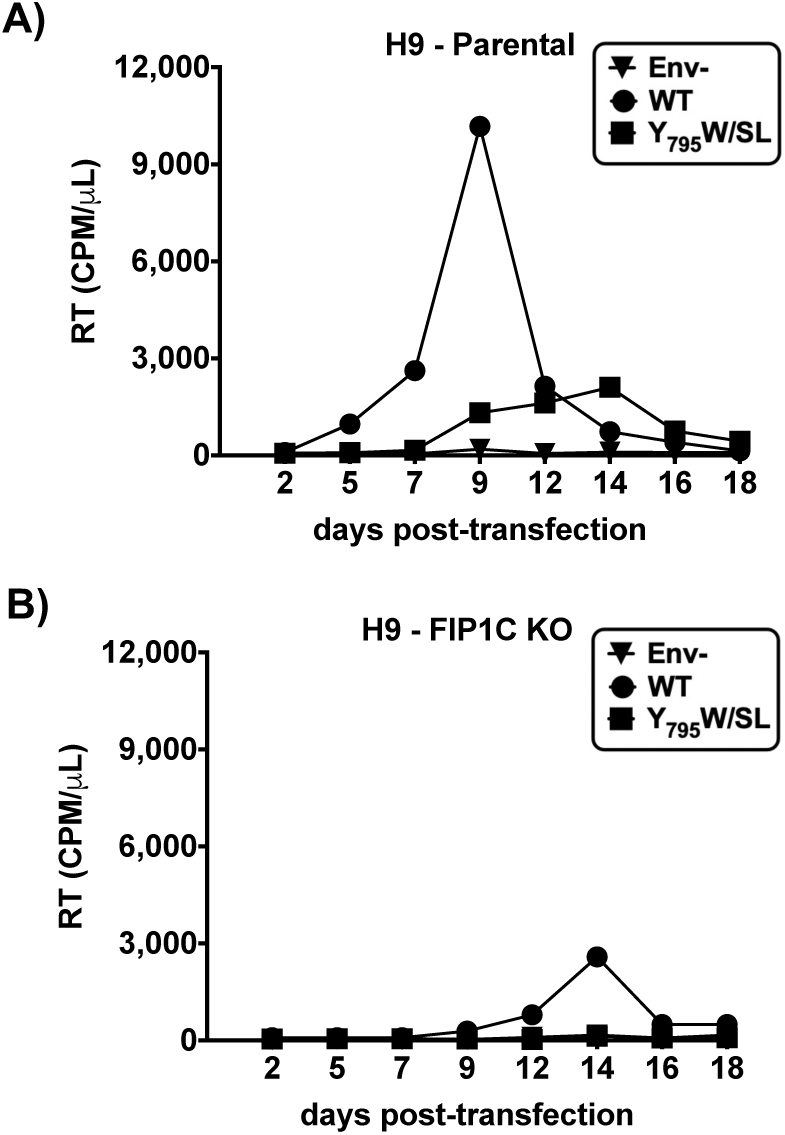
FIP1C KO in H9 cells results in attenuated virus replication. (**A–B**) Replication curves for spreading infection of WT and YW_795_SL NL4-3 in H9 and H9 FIP1C KO cells are shown. Cell lines were transfected with the proviral clone and split 1/3 and replenished with fresh RPMI-10% FBS every 2–3 days. An aliquot of the cell culture supernatant was reserved for analysis of HIV-1 RT activity at each time point. Data are representative of three independent experiments.

### Increasing cell density rescues detrimental effects of FIP1C KO on HIV-1 replication in H9 cells

A number of studies have demonstrated that, at least in cell culture, HIV-1 spreads much more efficiently via cell-cell than by cell-free transfer (see Introduction). We reasoned that increasing cell density would enhance the ability of the virus to spread via the cell-cell route. To examine this question, an equal number of cells were transfected with equivalent amounts of NL4-3 proviral DNA encoding either WT or Y_795_W/SL Env (Figure 8) and cultured at different densities by manipulating the culture surface area. Increasing the cell density of SupT1huR5 cultures resulted in a modest enhancement of the WT and Y_795_W/SL replication peak heights of both Parental and FIP1C KO cells (compare Figure 8A and B). In contrast, increasing the cell density of H9 cultures resulted in a complete rescue of HIV spreading infection kinetics in FIP1C KO cells (compare high density WT curves in Figure 8C and D). These data show that culturing cells at a high density can overcome the replication defect imposed by FIP1C KO in H9 cells.

**Figure 8.**
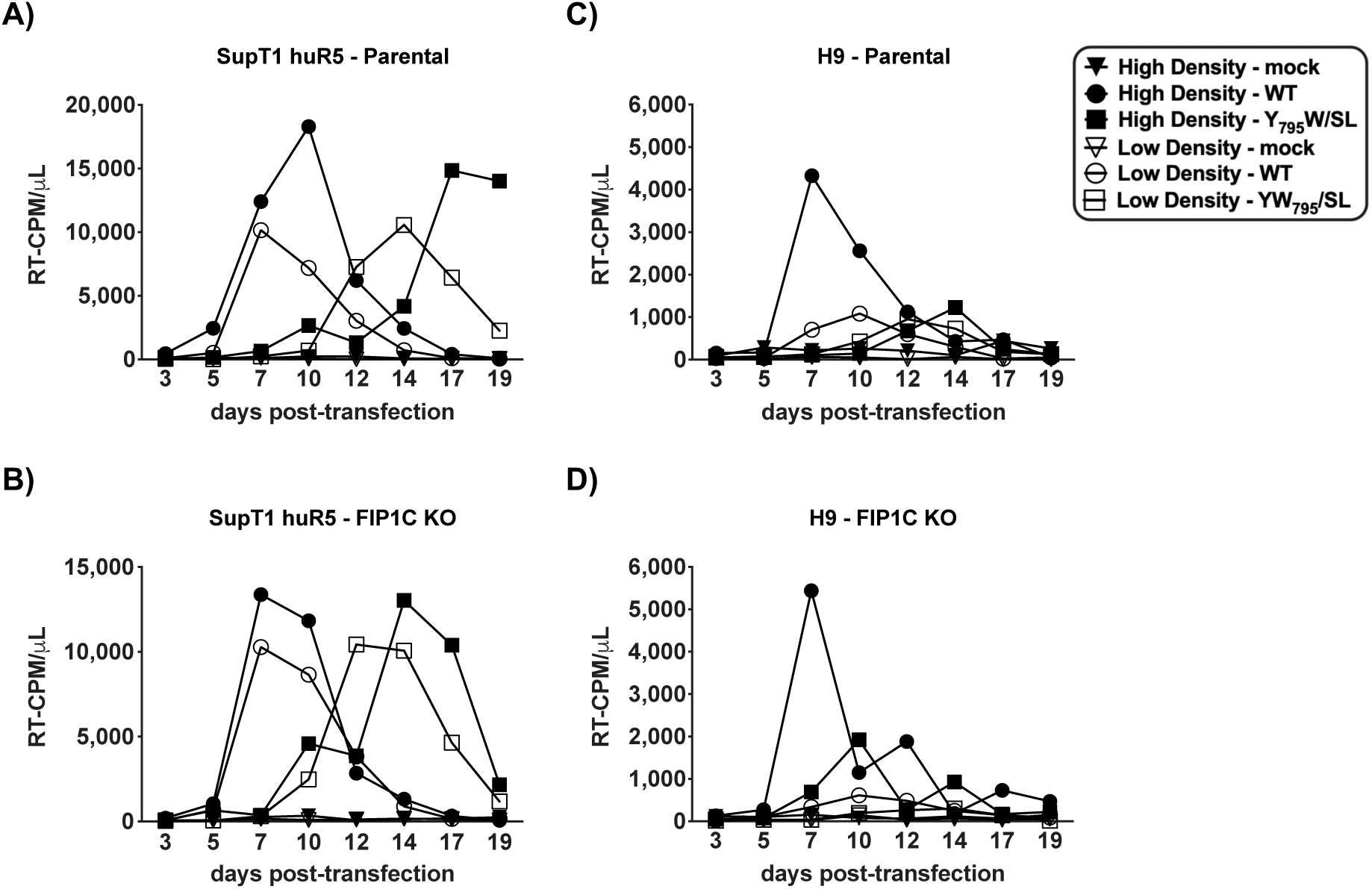
Increasing cell density rescues detrimental effects of FIP1C KO on HIV-1 replication in H9 cells. (**A–D**) Replication curves for spreading infection of WT and YW_795_SL NL4-3 in H9 and SupT1 huR5 parental and FIP1C KO cells are shown. Cells were transfected as described in the Materials and Methods. Cells were split 1/3 and replenished with fresh RPMI-10% FBS every 2– 3 days. An aliquot of the cell culture supernatant was reserved for analysis of HIV-1 RT activity at each time point. Data are representative of three independent experiments.

### Primary human PBMCs express FIP1C

To investigate whether FIP1C is expressed in primary human cells, and whether FIP1C expression is confined to CD4^+^ or CD4^−^ hPBMC subsets, three populations of PBMCs were generated: total, CD4^−^, and CD4^+^ cells. To generate the total hPBMC population, a fraction of the total PBMCs was separated and lysed. The remaining cells were used to generate the CD4^+^ and CD4^−^ cell populations. After CD4 enrichment and before lysing, a small portion of cells from the CD4-enriched and CD4-depleted fractions was reserved for cell-surface CD4 detection by flow cytometry. The results indicated that CD4 enrichment was highly efficient (Figure 9A). The resulting three populations were analyzed for FIP1C expression by western blotting using HeLa as a positive control for FIP1C expression (Figure 9B). The SupT1 T-cell line was included to allow a comparison between primary T cells and a T-cell line. FIP1C was expressed in all populations, albeit at different levels. FIP1C expression in SupT1 was higher than in HeLa or primary cells. Rab14 was also detected in HeLa, SupT1, CD4^−^, and CD4^+^ PBMCs.

**Figure 9.**
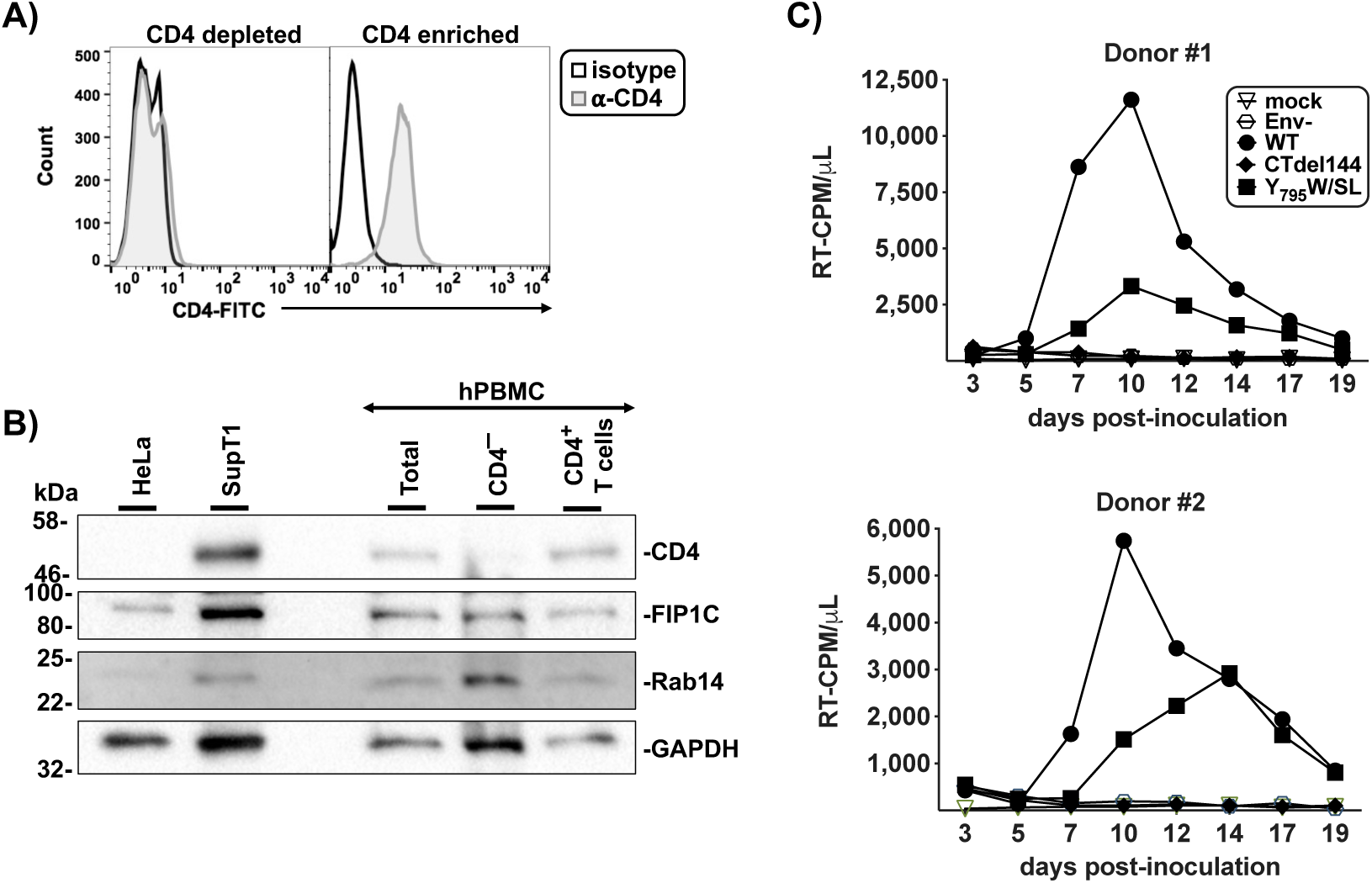
Primary human PBMCs express FIP1C. (**A**) CD4^+^ T cells were isolated from hPBMCs and analyzed by flow cytometry for CD4 expression. Data are representative of 3 donors. (**B**) CD4^−^ lymphocytes and CD4+ T cells were analyzed by Western blot analysis for FIP1C and Rab14, and compared to total PBMCs, and the FIP1C^+^ and Rab14^+^ SupT1 and HeLa cells. Data are representative of 3 donors. (**C**) hPBMCs were transduced with RT-normalized VSV-G pseudotyped NL4-3 encoding either Env(-), WT Env, CTdel144, or YW_795_SL Env. Samples were acquired as described in the Materials and Methods.

The results presented above and in a previous study (61) demonstrate that the Y_795_W/SL mutant is replication defective in T-cell lines. To determine whether this mutant can spread in primary CD4^+^ T cells, hPBMCs were transduced with VSV-G-pseudotyped NL4-3 encoding either Env(-), WT, CTdel144, or Y_795_W/SL Env. The virus-containing supernatant was sampled every 2–3 days to monitor virus replication. WT virus replication peaked 10 days post-inoculation while the Env(-) and CTdel144 Env mutants did not replicate, as expected (Figure 9C). The Y_795_W/SL Env mutant exhibited attenuated replication relative to WT, indicating that mutation of the Y_795_W motif results in impaired viral spread in primary CD4^+^ T cells.

### FIP1C is dispensable for NL4-3 replication in primary T cells

To determine if FIP1C is required for HIV-1 replication in primary CD4^+^ T cells, we used a CRISPR-Cas9 gene editing approach to knock-out FIP1C in cells from three independent healthy blood donors. Briefly, synthetic CRISPR guide RNAs (gRNA) were complexed *in vitro* with purified *S. pyogenes* Cas9 to form CRISPR-Cas9 ribonucleoprotein complexes, which were delivered to activated, primary CD4^+^ T cells by electroporation (Figure 10A) (71, 72). Four gRNAs targeting FIP1C were delivered independently as well as in a multiplexed pool alongside two non-targeting negative controls and previously validated gRNA targeting the known host and viral dependency factors CXCR4, CYPA, LEDGF, and TSG101 (71, 73). After allowing four days for DNA repair, protein depletion, and cell recovery, protein samples were harvested from each polyclonal culture for determination of knock-out efficiency. The guide RNA targeting FIP1C displayed varying efficiency with the multiplexed pool demonstrating near complete knock-out, similar to the CYPA control (Figure 10B). Cell viability was subsequently monitored by staining with a fluorescent amine dye and flow cytometry; no significant differences in cell viability were noted across each knock-out population (Figure 10C).

**Figure 10.**
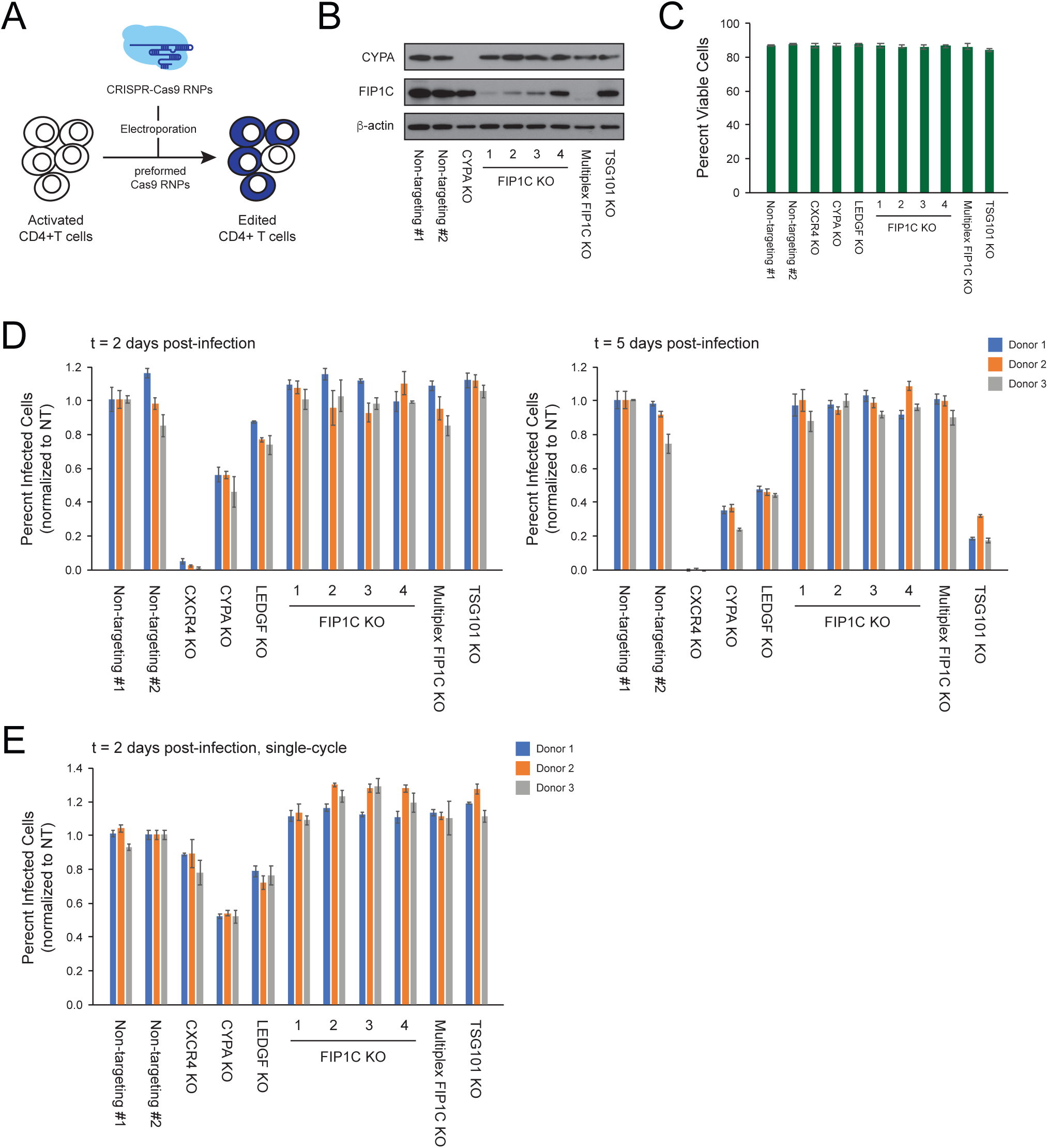
FIP1C is dispensable for NL4-3 replication in primary T cells. (**A**) Schematic of the CRISPR-Cas9 ribonucleoprotein knock-out approach used for this assay. (**B**) Immunoblots of each indicated knock-out population from one representative donor showing protein levels of CYPA and FIP1C relative to the b-actin loading control. (**C**) Percent viable CD4^+^ T cells in each indicated knock-out population as determined by amine dye staining and flow cytometry (mean ± standard deviation across three donors). (**D**) Percent infected (GFP+) cells 2-(left) and 5-days (right) post-challenge with HIV-1 NL4-3 nef:IRES:GFP in each indicated knock-out population (mean ± standard deviation across three technical replicates). Data are normalized to one of two non-targeting controls. (**E**) Percent infected (GFP+) cells 2-days post-challenge with VSV-G pseudotyped HIV-1 dEnv NL4-3 nef:IRES:GFP in each indicated knock-out population (mean ± standard deviation across three technical replicates). Data are normalized to one of two non-targeting controls.

Given the reported role of FIP1C in Env incorporation, we would not expect to see a replication phenotype in the knock-out population until the second round of viral replication. Each knock-out population was challenged in technical triplicate with replication-competent, CXCR4-tropic HIV-1 NL4-3 Nef:IRES:GFP. Infection rate (percent GFP+ cells) was monitored by flow cytometry at 2- and 5-days post-challenge and normalized to the first non-targeting control (Figure 10D). At day 2, there was almost no infection of the CXCR4 knock-out cells while infection of the CYPA and LEDGF knock-out cells was significantly decreased relative to the non-targeting controls. These differences were even more notable at day 5. The TSG101 knock-out cells showed no significant differences from the non-targeting controls at day 2, but had significantly less infection at day 5, consistent with the late-acting phenotype of TSG101.

None of the FIP1C knock-out cells showed any difference in infection relative to the non-targeting controls at either timepoint, regardless of gRNA efficiency. To test for the Env dependence of these phenotypes, these same cells were also infected in technical triplicate with single-cycle, VSV-G pseudotyped, HIV-1 dEnv NL4-3 Nef:IRES:GFP. Infection rate (percent GFP+ cells) was monitored by flow cytometry at 2-days post-challenge and normalized to the first non-targeting control (Figure 10E). As expected, VSV-G pseudotyping rescued infection of the CXCR4 knock-out cells while the CYPA and LEDGF knock-out cells showed similar decreases in infection to the replication-competent virus at day 2. The FIP1C and TSG101 knock-outs had no impact on infection of the single-round virus. Taken together, these data suggest that FIP1C is dispensable for HIV-1 NL4-3 replication in activated, primary CD4^+^ T cells.

## Discussion

The function of the long gp41 CT of primate lentiviruses, and the identity and role of host factors that interact with the gp41 CT, has long been enigmatic. It is well established that clathrin adapter protein complexes bind the gp41 CT and direct Env internalization from the plasma membrane (4–10). Other factors, including retromer (74), have been demonstrated to interact with the gp41 CT [for review, see P. R. Tedbury and E. O. Freed (3)]; however, the role that CT-interacting factors play in Env incorporation and virus propagation remains incompletely understood. To broadly evaluate the role of FIP1C in HIV-1 Env incorporation, particle infectivity, and virus spread, we knocked out FIP1C expression in multiple T-cell lines and primary T cells by using CRISPR/Cas9 technology. We observed a range of phenotypes resulting from *FIP1C* KO. In the Jurkat T-cell line, Env incorporation or virus replication was not reduced. In the SupT1 T-cell line, modest (∼33%) reductions in Env incorporation efficiency and single-cycle infectivity were observed, but virus replication kinetics were not affected by *FIP1C* KO. In H9 cells, we observed delayed virus replication, consistent with the results of Qi et al. (60). In this T-cell line, *FIP1C* KO caused a ∼33% reduction in Env incorporation efficiency and an overall reduction in Env levels, suggesting that FIP1C KO may reduce the stability of Env in this cell line, perhaps by disrupting Env trafficking. Interestingly, H9 expressed more FIP1C protein than SupT1 and were the only cells tested in this study in which FIP1C KO attenuated WT NL4-3 replication. The virus replication defect observed in *FIP1C* KO H9 cells could be rescued by increasing cell density. Because higher cell density and the corresponding increase in cell-cell contacts favor cell-cell virus transmission, these results suggest that the cell-cell mode of HIV-1 spread may be less susceptible to the loss of FIP1C expression than the cell-free mode. Indeed, cell-cell transfer has been reported to enable HIV-1 to overcome a number of blocks to virus replication (28–32). Perhaps most importantly, *FIP1C* KO did not detectably impair HIV-1 replication in primary CD4^+^ T cells.

We examined Env incorporation and virus replication in two single-cell-derived *FIP1C* KO Jurkat E6.1 clones, denoted clone #6 and #9. In clone #6, we observed no significant effect of *FIP1C* KO on Env incorporation, particle infectivity, or virus replication. This result indicates that FIP1C is not required for HIV-1 replication in the Jurkat E6.1 T-cell line. In clone #9, we observed an increase in viral protein (Env and Gag) expression, and, correspondingly, more rapid HIV-1 replication kinetics relative to parental cells. In our view, considering the results obtained with clone #6, it is unlikely that the increases in viral gene expression and replication kinetics observed with clone #9 were due to the *FIP1C* KO but rather were probably a property of the single cell from which this clone was derived. These results underscore the importance of analyzing multiple single-cell-derived clones knocked out for a gene of interest or using polyclonal KO cell lines that avoid artifacts stemming from a single-cell origin.

Also underscoring the need to evaluate many cell lines when evaluating novel host factors in viral assembly/replication, in addition to primary T cells, is the finding that many HTLV-I transformed cells (specifically ED) do not express FIP1C. ED was not permissive to CTdel144 replication but was permissive to WT HIV-1 replication (data not shown), indicating that they do not behave like MT-4 cells; rather, they behave like non-HTLV-1 transformed T-cell lines in their ability to support HIV-1 replication. Therefore, these data indicate that FIP1C either does not impose a restriction on plasma membrane Env levels or its deficiency is compensated by a redundant trafficking protein.

As described in the introduction, mutation of the Y_795_W motif in the gp41 CT has been shown to play a role in HIV-1 Env incorporation and particle infectivity (37, 39) and this motif has been implicated in FIP1C-dependent Env incorporation and virus replication (61, 62). The Y_795_W motif was also reported to be required for the Env-dependent relocalization of FIP1C from a perinuclear compartment (potentially a recycling endosome) to the plasma membrane, suggesting that this motif might be required for direct or indirect interactions between gp41 and FIP1C (62).

In this study, we confirmed that the Y_795_W/SL mutation reduces Env incorporation and impairs HIV-1 replication across a range of cell lines. We also demonstrated a role for the highly conserved Tyr residue in this motif in the replication of subtype C HIV-1. However, in our experiments, the defects exhibited by the Y_795_W/SL NL4-3 mutant, and the Y_795_S subtype C mutant, were independent of FIP1C expression, as similar phenotypes for these mutants were observed in parental and *FIP1C* KO cells. An NMR solution structure of the gp41 CT embedded in lipid micelles suggested that a region at the N-terminus of LLP-3 that is rich in aromatic residues, including Y_795_W, interacts closely with the inner leaflet of the membrane (50). The Y_795_W motif also lies within, and adjacent to, regions of the gp41 CT that have been proposed to interact with MA (37, 51, 75). Mutation of the Y_795_W motif could thus affect the mobility of Env in the membrane, local Env conformation, or interaction of the gp41 CT with the MA domain of Gag, thereby disrupting Env incorporation. Interestingly, Qi et al. (61) reported that the Y_795_W/SL mutant reverted via acquisition of a single amino acid mutation (L_850_/S) near the C-terminus of the gp41 CT. It remains to be determined by what mechanism this second-site change rescues the defects exhibited by the Y_795_W/SL mutant.

In summary, the results presented here indicate that while FIP1C appears to play a modest role in Env incorporation in some cell lines, it is not required for HIV-1 replication in most T-cell lines tested, or in primary CD4^+^ T cells. Further studies will continue to clarify the role of the enigmatic gp41 CT in HIV-1 replication and its interaction with host cell trafficking machinery.

## Materials and Methods

### Cell lines and tissue culture

HeLa (a gift from P. Spearman), 293T [obtained from American Type Culture Collection (ATCC)] and TZM-bl [obtained from J. C. Kappes, X. Wu, and Tranzyme, Inc. through the NIH AIDS Reagent Program (ARP), Germantown, MD] cell lines were maintained in DMEM containing 5% or 10% (vol/vol) fetal bovine serum (FBS), 2 mM glutamine, 100 U/mL penicillin, and 100 µg/mL streptomycin (Gibco) at 37°C with 5% CO_2_. Jurkat E6.1 [for reference, see (64); Cat# ARP-177; cell accession ID on the Cellosaurus database is CVCL_0367], MT-4 [STR profile published in (64, 66); Cat# ARP-120; cell line accession ID on the Cellosaurus database is CVCL_2632), C8166-45 (referred to as C8166; cell line accession ID on the Cellosaurus database is CVCL_0195; Cat# ARP-404), and M8166 [for reference, see (64); Cat# ARP-11395; cell line accession ID on the Cellosaurus database is CVCL_1H07], were obtained from Arthur Weiss, Douglas Richman, Robert Gallo, and Paul Clapham, respectively, through the NIH ARP. The laboratory source of SupT1 used in this study is unknown [for reference, see (64)]. H9 cells were obtained from Robert Gallo (Cat# ARP-87) through the NIH-ARP and STR profiled to confirm their identity and was a 100% match to the Cellosaurus cell line accession ID CVCL_1240. The SupT1-CCR5 (76) (referred to as SupT1huR5; cell line accession ID on the Cellosaurus database is CVCL_X633) T-cell line was a generous gift from James Hoxie [cell line accession ID on the Cellosaurus database is CVCL_X633; STR profile published in (64)]. The ATL Tax^−^ cells lines (ED, ATL-55T, and TL-Om1) were a generous gift from Chou-Zen Giam. T-cell lines were maintained in Roswell Park Memorial Institute (RPMI)-1640 medium containing 10% FBS, 2 mM glutamine, 100 U/mL penicillin, and 100 µg/mL streptomycin (Gibco) at 37°C with 5% CO_2_. Whole blood was obtained from healthy donors via the NCI-Frederick Research Donor Program. hPBMCs were isolated using a ficoll gradient (Histopaque-1077; Sigma #10771) and stimulated with 2 μg/mL phytohemagglutinin P (PHA-P) for 3–5 days before infection, then cultured in 50 U/mL IL-2.

### Cloning and plasmids

Unless otherwise indicated, the full-length HIV-1 subtype B molecular clone pNL4-3 was used (77). The Env-deficient pNL4-3/KFS clone, referred to here as Env(−) (78) and gp41 CT truncation mutant CTdel144 (36) were described previously. The pNL4-3 Y_795_W/SL molecular clone was generated using the QuikChange Site-Directed Mutagenesis Kit (Agilent Technologies) on pSL1190-EcoRI-Env WT-XhoI, and cloned into pNL4-3 by EcoRI and XhoI restriction digest of pSL1190-EcoRI-Env Y_795_W/SL-XhoI and pNL4-3, followed by ligation of the *env* fragment into pNL4-3. The primers were designed to retain the overlapping Rev amino acid sequence. Plasmid sequences were confirmed by restriction digest with HindIII and Sanger sequencing.

Primers used for mutagenesis are as follows:

NL4-3 Y_795_W/SL FWD 5’-GGGGTGGGAAGCCCTCAAATCCTTGTGGAATCTCCTAC-3’ NL4-3 Y_795_W/SL REV 5’-GTAGGAGATTCCACAAGGATTTGAGGGCTTCCCACCCC-3’ The full-length HIV-1 transmitted/founder subtype C molecular clones pK3016 (CH185) and pK3058 (CH067) were reported previously (79) and were a gift from C. Ochsenbauer. The pK3016 Y_795_S molecular clone was engineered with the Q5 Mutagenesis Kit (New England Biolabs) according to manufacturer’s instructions. The primers used did not preserve the overlapping Rev amino acid sequence, but the mutations did not affect viral protein expression, indicating no defect in Rev function. pK3016 and pK3058 were amplified using Stbl3 cells (Thermofisher) and standard LB medium. DNA for transfections was purified in large-scale quantities using MaxiPrep Kits (Qiagen) and mutations were verified by Sanger sequencing (Macrogen). Plasmid sequences were confirmed by restriction digest with HindIII and Sanger sequencing of the full plasmid.

Primers used for mutagenesis are as follows:

K3016 Y_795_S FWD: GGGGGTGGGAAGCCCTGAAGTCCCTGGGAAGTCTTGTGC

K3016 Y_795_S REV: GCACAAGACTTCCCAGGGACTTCAGGGCTTCCCACCCCC

### siRNA-mediated knockdown of FIP1C

*FIP1C* knockdown in HeLa cells was accomplished by reverse siRNA transfection. Briefly, 100 pmol of siRNA was diluted in 500 uL of Opti-MEM in a well of a 6-well plate and gently mixed. 5 uL of Lipofectamine RNAiMax (Thermofisher) was added to each well and mixed gently with the siRNA and incubated for 15 min at room temperature. HeLa cells were diluted in DMEM without serum at a concentration of 2×10^5^ cells/mL. 0.7 mL of the diluted HeLa cells were added to the siRNA transfection complex and mixed. The cells were mixed gently then incubated at 37°C for 4 h. After incubation, 1 mL of DMEM-5% FBS without antibiotics was added to each well. Cells were incubated for 48 h at 37°C and then transfected with pNL4-3 by using Lipofectamine 2000 (Thermofisher) according to the manufacturer’s instructions. Transfected cells were incubated for 2 days before viral and cell lysates were collected.

### Preparation of virus stocks

293T cells were transfected with HIV-1 proviral DNA using Lipofectamine 2000 (Invitrogen) according to the manufacturer’s instructions. Virus-containing supernatants were filtered through a 0.45 µm membrane 48 h post-transfection and virus was quantified by measuring RT activity. VSV-G–pseudotyped virus stocks were generated from 293T cells co-transfected with proviral DNA and the VSV-G expression vector pHCMV-G (80) at a DNA ratio of 10:1.

### STR profiling and cell line validation

The identity of T-cell lines was confirmed by performing STR profiling as described previously (66). Briefly, genomic DNA was extracted and sent to Genetica (LabCorp) for profiling. The URL for this data set is: https://amp.pharm.mssm.edu/Harmonizome/gene_set/IRF3/ENCODE+Transcription+Factor+Targets. The obtained STR profile was compared to the Cellosaurus reference STR using the % Match formula as previously described (64).

### Illumina RNA-Seq

RNA was extracted from T cells in their exponential growth phase using QIAshredder (Qiagen, Cat# 79654) and RNeasy Plus Mini Kit (Qiagen, Cat# 74134). RNA sample integrity was assessed by determining the RNA Integrity Number (RIN) with an Agilent 2100 Bioanalyzer instrument and applying the Eukaryote Total RNA Nano assay. RIN values were between 9 and 10, indicating intact RNA. Samples were sequenced on a Hiseq 2500, generating an average of 50 million raw reads per sample. Transcript sequence reads were normalized against the total reads for each cell line to generate a reads per kilobase per million mapped reads (RPKM) value. The RPKM is a relative measure of transcript abundance.

### Virus Replication Assays

Virus replication kinetics were determined as previously described (64). Briefly, 2.5×10^6^ T cells were transfected with proviral clones (1 μg DNA/10^6^ cells) in the presence of 700 μg/mL DEAE-dextran and cultured in 3 mL of RPMI-10% FBS. Every 2–3 days a 200 μL sample was collected from each flask for RT assay and frozen in a 96-well plate before splitting the cell cultures (Jurkat E6.1, SupT1, SupT1huR5, and H9) 1:3 with fresh media.

To test the effect of varying cell density on replication kinetics, an equivalent number of cells (2.5×10^6^) were transfected with 0.1 μg DNA/10^6^ cells in duplicate and maintained in 5 mL of RPMI-10% FBS. To alter the cell density, one replicate was maintained in a T-25 cm^2^ flask upright (high cell density) and the other replicate was maintained with a T-25 cm^2^ flask lying flat (low cell density).

VSV-G-pseudotyped virus was used to inoculate PHA-stimulated hPBMCs. Virus stocks were normalized by RT activity and used to initiate spreading infection. After a 2 h incubation with VSV-G-pseudotyped virus, cells were washed and resuspended in fresh RPMI-10% FBS +50 Units rhIL-2. Every other day, half the medium was replaced without disturbing the cells. Virus replication was monitored by measuring the RT activity in collected supernatants over time. RT activity values were plotted using GraphPad Prism to generate replication curves.

### HIV-1 infection of T cells and generation of viral and cell samples for analysis of Env content

293T cells were plated and co-transfected via lipofectamine the next day with the pNL4-3 proviral clone and the VSV-G expression vector, pHCMV-G, at a 10:1 ratio. 48 h post-transfection, supernatants were passed through a 0.45 µm filter and RT activity was measured as described (81). T cells were plated the night before infection in fresh media at a density of 5×10^6^ cells / 2 mL. The following day, cells were infected overnight with RT-normalized virus. After an overnight incubation with virus, cells were washed extensively and cultured in 1.5 mL of RPMI-10% FBS. The virus and cell fractions were collected 40 h post-transduction for SupT1huR5 and Jurkat E6.1 and 72 h post-transduction for H9. H9 were spinoculated for 2 h at 25°C and 200 x *g* to initiate the transduction while Jurkat E6.1 and SupT1huR5 were not spinoculated.

Virus-containing supernatants were passed through a 0.45µm filter. 10 µL were set aside for RT assay and 200 µL were set aside for TZM-bl infectivity assay. The remaining filtered virus-containing supernatant was layered on a 20% w/v sucrose/phosphate buffered saline (PBS) solution and spun for 1.25 h at 100,000 x g at 4°C in a Sorvall S55-A2 fixed angle rotor (ThermoFisher Scientific). Cell and virus fractions were solubilized in lysis buffer [30 mM NaCl, 50 mM Tris-HCL pH 7.5, 0.5 % Triton X-100, 10mM Iodoacetamide, complete protease inhibitor (Roche)].

### Protein detection

For chemiluminescence-based detection of viral proteins, lysates boiled with 6x loading buffer (7 mL 0.5 M Tris-HCL/0.4 % SDS, 3.8 g glycerol, 1 g SDS, 0.93 g DTT, 1.2 mg bromophenol blue) were subjected to SDS-PAGE on 12% 1.5 mm gels and processed using standard western blotting techniques. All antibodies were diluted in 10 mL of 5% milk in TBS blocking buffer. HIV proteins were detected with 10 µg/mL polyclonal HIV immunoglobulin (HIV-Ig) obtained from the NIH ARP. Anti-human IgG conjugated to horseradish peroxidase [HRP; SigmaAldrich (Cat# GENA933)] and used at a 1:5,000 dilution. gp41 was detected with 2 µg/mL10E8 monoclonal antibody obtained from the NIH ARP (Cat# 12294) followed by anti-human IgG-HRP as above. Protein bands were visualized using chemiluminescence with a Bio-Rad Universal Hood II Chemidoc and then analyzed with ImageLab v5.1 software.

For fluorescence-based detection of viral proteins, cell lysates were prepared as above and loaded on 4–15% pre-cast gels (BioRad). After protein separation, proteins were transferred with the trans-blot turbo transfer system (BioRad) according to the manufacturer’s instructions using the manufacturer’s preset 30 min Standard SD protocol onto a low-fluorescence PVDF membrane (Thermofisher). After the transfer, the membrane was blocked for 30 min in 10 mL azure fluorescent blot blocking buffer (Azure Biosystems). The antibodies 16H3, chessie8, and HIV-Ig were diluted 1/5,000 all together in azure fluorescent blot blocking buffer and 10 mL of the diluted antibodies were added to the membrane. The membrane was incubated with antibodies at 4°C overnight, washed according to the manufacturer’s instructions with azure fluorescent blot washing buffer (Azure Biosystems) according to the manufacturer’s instructions, and incubated with azure spectra fluorescent secondary antibodies goat-anti-mouse 800 and goat-anti-human 650 at 4°C overnight. The following day the membrane was washed extensively with azure fluorescent blot washing buffer followed by a quick wash with methanol, and then dried. The membrane was then scanned with the azure biosystems sapphire biomolecular imager and analyzed with the azure spot analysis software.

For detection of cellular proteins, cell lysates were processed as described above and incubated with the following antibodies: anti-GAPDH (1:20,000 dilution; Abcam, catalog #ab26997), anti-β-actin conjugated directly to horseradish peroxidase (HRP; Abcam, catalog #ab49900), anti-Rab14 (Abcam, catalog #ab28639), and anti-FIP1C (Cell Signalling catalog #12849). Protein bands were visualized using chemiluminescence with a Bio-Rad universal hood II chemidoc and then analyzed with ImageLab v5.1 software or with the azure biosystems sapphire biomolecular imager and then analyzed with ImageLab v5.1 software or the azure spot analysis software.

### CD4^+^ cell enrichment for FIP1C analysis

Human buffy coats were obtained from healthy volunteers through the NCI-Frederick Research Donor Program. PBMCs were isolated by ficoll gradient as reported (82). CD4^+^ T cells were isolated from PBMCs by magnetic column isolation using the CD4+ T-cell isolation kit (Miltenyi Biotec catalog #130-096-533). CD4^-^ cells were also collected.

### Single-cycle infectivity assays

TZM-bl infectivity assays were performed as previously described (31). Briefly, 20,000 TZM-bl cells were plated in a flat-bottomed, white-walled plate (Sigma Aldrich, Cat# CLS3903-100EA). The following day, the cells were infected with serial dilutions of RT-normalized virus stocks in the presence of 10 µg/mL DEAE-dextran. Approximately 36 h post-infection, cells were lysed with BriteLite luciferase reagent (Perkin-Elmer) and luciferase was measured in a Wallac 1450 Microbeta Counter plate reader.

### Generation of stable FIP1C KO Jurkat E6.1, SupT1 HuR5, and H9

The LentiCRISPRv2 plasmid, which codes for Cas9 endonuclease and a puromycin resistance selection marker as well as containing a cloning site for sgRNA expression, was a gift from Feng Zhang (Addgene plasmid #52961). A single guide RNA sequence (sgRNA) targeting a site within the first exon of FIP1C was selected by using the CRISPOR design program (83) to identify sgRNAs with high specificity, efficiency, and likelihood of producing out-of-frame indels. The chosen FIP1C sgRNA target sequence was GGGCACGAGCGACGCGTACG. The sgRNA was cloned into LentiCRISPRv2 as previously described (84), to generate the transfer plasmid LentiCRISPRv2-FIP1CsgRNA.

Lentivirus was produced by transfecting HEK293T cells with the LentiCRISPRv2-FIP1CsgRNA transfer plasmid along with the psPAX2 packaging plasmid (gift from Didier Trono, Addgene plasmid #12260) and pVSV-G pseudotyping plasmid (gift from Dr. Xuedong Liu, University of Colorado, Boulder), using PEI transfection reagent (Alfa Aesar/Thermo Fisher Scientific, Tewksbury, MA, #43896).

Jurkat E6.1, H9, and SupT1huR5 cells were transduced with LentiCRISPRv2-FIP1CsgRNA and selected with 500 ng/mL puromycin beginning two days after transduction. After puromycin selection was complete, clonal lines of FIP1C KO Jurkat E6.1 cells were isolated by limiting dilution. FIP1C KO H9 and FIP1C KO SupT1huR5 cells were used without clonal isolation.

### CRISPR-Cas9 RNP production, CD4^+^ T cell isolation, and electroporation of primary CD4^+^ T cell cultures

Detailed protocols for RNP production and primary CD4+ T cell editing have been previously published (72). Briefly, lyophilized CRISPR-Cas9 ribonucleoproteins (crRNA) and trans-activating CRISPR RNA (tracrRNA) (Dharmacon) were suspended at a concentration of 160 µM in 10 mM Tris-HCl (7.4 pH) with 150 mM KCl. 5 µL of 160 µM crRNA was mixed with 5µL of 160µM tracrRNA and incubated for 30 min at 37°C. The crRNA:tracrRNA complexes were then mixed gently with 10 µL of 40 µM Cas9 (UC-Berkeley Macrolab) to form Cas9 ribonucleoproteins (RNPs). Five 3.5 µL aliquots were frozen in Lo-Bind 96-well V-bottom plates (E&K Scientific) at -80°C until use. For multiplex RNP synthesis, multiple crRNA targeting the same gene were mixed in equal ratios before proceeding with tracrRNA addition as above. All crRNAs used in this study were purchased as custom sequences from Dharmacon or derived from the Dharmacon pre-designed Edit-R library for gene knock-out (see crRNA sequences table below).

Primary human CD4^+^ T cells from healthy donors were isolated from leukoreduction chambers after Trima Apheresis (StemCell). PBMCs were isolated by Ficoll centrifugation. Bulk CD4^+^ T cells were subsequently isolated from PBMCs by magnetic negative selection using an EasySep Human CD4^+^ T Cell Isolation Kit (STEMCELL, per manufacturer’s instructions). Isolated CD4^+^ T cells were suspended in RPMI-1640 (Sigma) supplemented with 5 mM 4-(2-hydroxyethyl)-1-piperazineethanesulfonic acid (HEPES, Corning), 50 μg/mL penicillin/ streptomycin (P/S, Corning), 5mM sodium pyruvate (Corning), and 10% FBS (Gibco). Media were supplemented with 20 IU/mL IL-2 (Miltenyi) immediately before use. For activation, bulk CD4^+^ T cells were immediately plated on anti-CD3 coated plates (coated for 2 h at 37°C with 20 µg/mL anti-CD3 [UCHT1, Tonbo Biosciences]) in the presence of 5 µg/mL soluble anti-CD28 (CD28.2, Tonbo Biosciences). Cells were stimulated for 72 h at 37°C / 5% CO_2_ prior to electroporation.

Each electroporation reaction consisted of 5×10^5^ activated T cells, 3.5 µL RNPs, and 20 µL electroporation buffer. After three days of stimulation, cells were suspended and counted. RNPs were thawed and allowed to warm to room-temperature. Immediately prior to electroporation, cells were centrifuged at 400xg for 5 min, supernatant was removed by aspiration, and the pellet was resuspended in 20 µL of room-temperature P3 electroporation buffer (Lonza) per reaction. 20 µL of cell suspension was then gently mixed with each RNP and aliquoted into a 96-well electroporation cuvette for nucleofection with the 4D 96-well shuttle unit (Lonza) using pulse code EH-115. Immediately after electroporation, 80 µL of pre-warmed media with IL-2 were added to each well and cells were allowed to rest for at least 1 h in a 37°C cell culture incubator. Cells were subsequently moved to 96-well flat-bottomed culture plates pre-filled with 100 µL warm complete media with IL-2 and anti-CD3/anti-CD2/anti-CD28 beads (T cell Activation and Stimulation Kit, Miltenyi) at a 1:1 bead:cell ratio. Cells were cultured at 37°C / 5% CO_2_ in a humidified cell culture incubator for 4 days to allow for gene knock-out and protein clearance, with additional media added on day 2. To collect protein lysates for determination of knock-out efficiency, 50 µL of each mixed culture was removed to a centrifuge tube. Cells were pelleted, supernatant was removed, and pellets were resuspended in 75 µL 2.5x Laemmli Sample Buffer. Protein lysates were heated to 98°C for 20 min before storage at -20°C until immunoblotting.

### crRNA Sequences

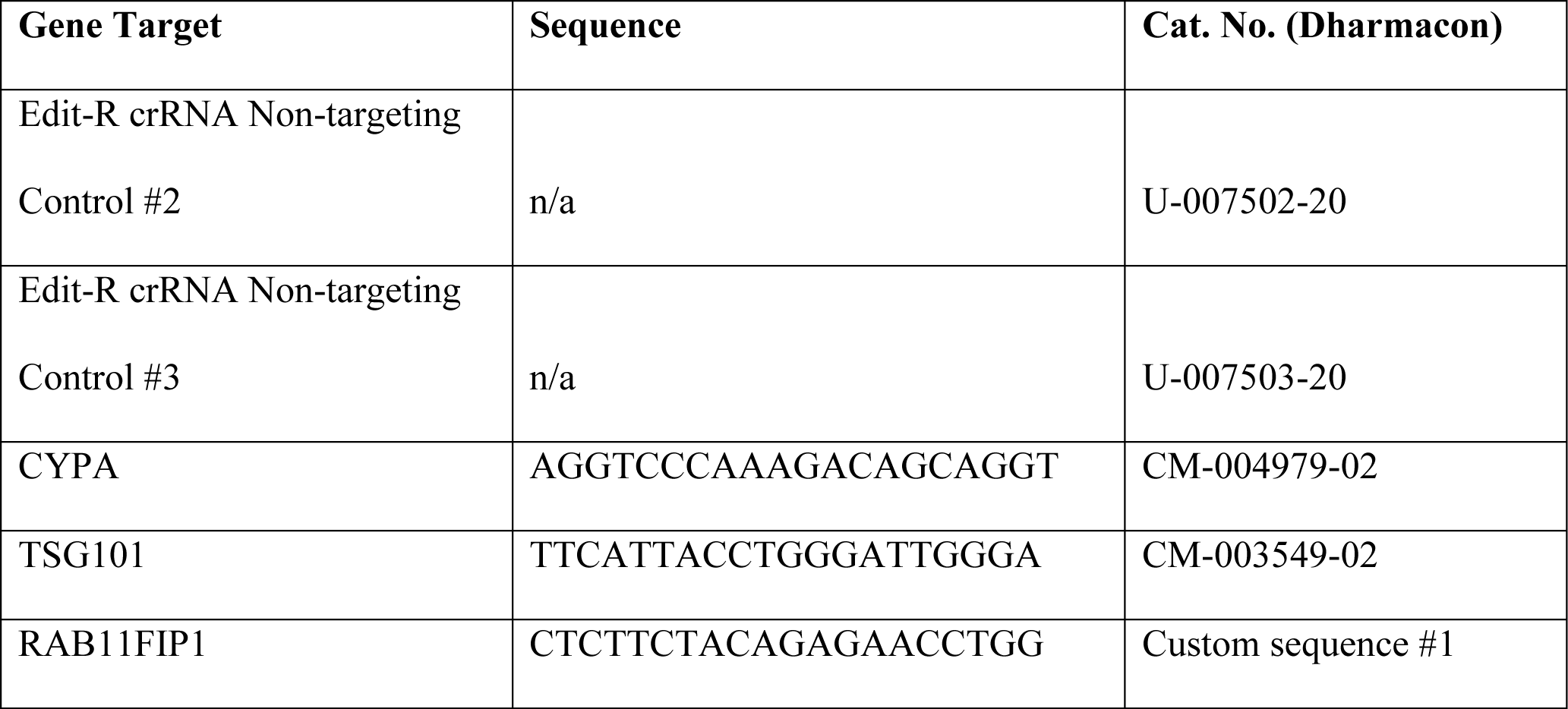
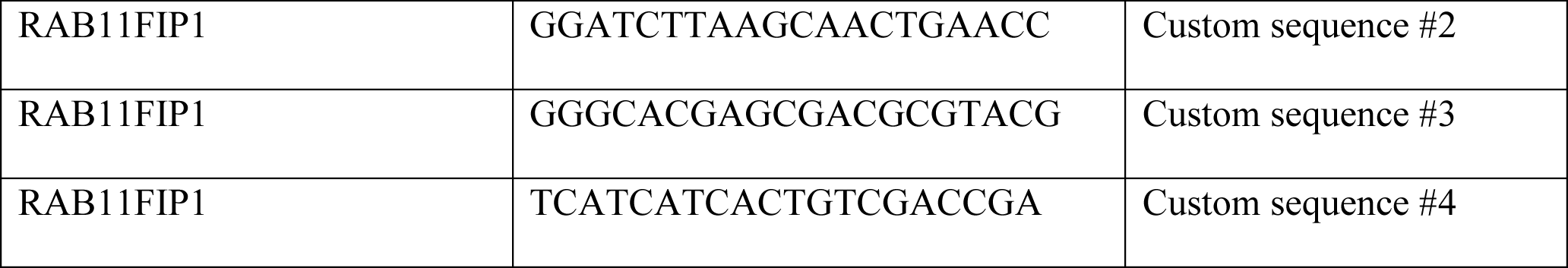

### Immunoblotting of KO primary CD4^+^ T cell cultures

Samples and PageRuler™ Plus Prestained Protein Ladder were thawed and 15 µL of each was loaded onto 4-20% Criterion Tris-HCl SDS-PAGE protein gels (BioRad). Gels were run at 150V over 90 min until the ladder was sufficiently separated. Proteins were transferred to PVDF membranes by methanol-based electrotransfer (BioRad Criterion Blotter) at 90V for 2 h. Membranes were blocked in 4% Milk in PBS, 0.1% Tween-20 overnight prior to overnight incubation with primary antibody against CYPA (clone 79D7, Cell Signaling Technologies), RAB11FIP1 (clone D9D8P, Cell Signaling Technologies), and 1-actin (clone 8H10D10, Cell Signaling Technologies) as a protein loading control. Anti-rabbit or anti-mouse horseradish peroxidase (HRP)-conjugated secondary antibodies (BioRad) were detected using Hyglo HRP detection reagents (Denville Scientific). Blots were incubated in a 1xPBS, 0.2M glycine, 1.0% SDS, 1.0% Tween-20, pH 2.2 stripping buffer before reprobing.

### Preparation of virus stocks for infection of primary CD4^+^ T cell cultures

Replication-competent reporter virus stocks were generated from an HIV-1 NL4-3 molecular clone in which GFP had been cloned behind an internal ribosomal entry site (IRES) cassette following the viral *nef* gene (NIH AIDS Reagent Program, cat. no. 11349). Briefly, 10 µg of molecular clone was transfected (PolyJet, SignaGen) into 5×10^6^ HEK293T cells (ATCC CRL-3216) according to the manufacturer’s protocol. 25 mL supernatant was collected at 48 and 72 h and combined. Virus-containing supernatant was filtered through 0.45 mm PVDF filters (Millipore) and precipitated in 8.5% polyethylene glycol (PEG, average M_n_ 6000, Sigma), 0.3 M NaCl for 4 h at 4°C. Supernatants were centrifuged at 3500 rpm for 20 min and virus resuspended in 0.5 mL phosphate buffered saline (PBS) for a 100x effective concentration.

Aliquots were stored at -80°C until use. Single-round reporter virus stocks were generated from a similar HIV-1 NL4-3 dEnv molecular clone with an integrated nef:IRES:GFP. Viral particles were pseudotyped with VSV-G by co-transfection with pMD2.G (Addgene cat. co. 12259) at a 3:1 ratio of proviral clone:pMD2.G prior to purification as above.

### HIV-1 infection of primary CD4^+^ T cell cultures

Detailed protocols for HIV-1 spreading infection have been previously described (72). Briefly, 6 days post-electroporation, cells were replica-plated into triplicate, 96-well, round-bottom plates and cultured overnight in 150 µL complete RPMI as described above in the presence of 20 IU/mL IL-2. On the following day, 2.5 µL of concentrated virus stock was added to each well in a 50 µL carrier volume to bring the total volume in each well to 200 µL. Cells were cultured in a dark, humidified incubator at 37°C / 5% CO_2_. On days 2 and 5 post-infection, 75 µL of each culture was removed and mixed 1:1 with freshly made 2% formaldehyde in PBS (Sigma) and stored at 4°C for analysis by flow cytometry. Cultures were supplemented with 75 µL complete, IL-2-containing RPMI media and returned to the incubator. For single-round infection assays, cells were plated and challenged with 2.5 µL of single-round, concentrated virus stock as above. Two days post-challenge, 75 µL of each culture was removed and mixed 1:1 with freshly made 2% formaldehyde in PBS (Sigma) and stored at 4°C for analysis by flow cytometry.

### Flow cytometry and analysis of primary CD4^+^ T cell infection data

Flow cytometric analysis was performed on an Attune NxT Acoustic Focusing Cytometer (ThermoFisher), recording all events in a 100 µL sample volume after one 150 µL mixing cycle. Data were exported as FCS3.0 files and analyzed with a consistent template on FlowJo. Briefly, cells were gated for lymphocytes by light scatter followed by doublet discrimination in both side and forward scatter. Cells with equal fluorescence in the BL-1 (GFP) channel and VL-2 (AmCyan) channels were identified as auto-fluorescent and excluded from analysis. A consistent gate was then used to quantify the fraction of remaining cells that expressed GFP.

### Statistics

Statistics were calculated using GraphPad Prism version 8 for Mac OS (GraphPad Software, La Jolla, CA). Unpaired Student’s t-tests were performed and two-tailed *P < 0.05, **P < 0.01, ***P < 0.001, and ****P < 0.0001 were considered statistically significant. GraphPad Prism was also used to calculate standard error and to assess statistical significance by one-way ANOVA. P values for Student’s t-test and one-way ANOVA analysis are defined with the same cut-offs.

### Ethics Statement

PBMCs were obtained from anonymous, de-identified blood donors through the NCI-Frederick Research Donor Program or were purchased through StemCell.

## List of abbreviations

CT: cytoplasmic tail
VS: virological synapse
C-C: cell-to-cell
Env: envelope glycoprotein
STR: short tandem repeat
PM: plasma membrane
hPBMC: human peripheral blood mononuclear cells
RT: reverse transcriptase
VSV-G: Vesicular stomatitis virus glycoprotein
PBS: phosphate buffered saline
FBS: fetal bovine serum
WT: wild type
HRP: horseradish peroxidase
MA: HIV-1 Matrix protein
NIH ARP: NIH AIDS Reagent Program
n.s.: not statistically significant
CA: HIV-1 Capsid protein
HTLV-I: human T-cell leukemia virus type I
ATL: adult T-cell leukemia/lymphoma
RIN: RNA Integrity Number

## Acknowledgments

We thank Paul Spearman for providing the HeLa cells used in this study, James Hoxie for providing the SupT1huCCR5 cell line, and Chou-Zen Giam for helpful discussion and for providing the ATL Tax– T cells (ED, ATL-55T, and TL-Om1). We also thank Yongmei Zhao at the NCI-Center for Cancer Research Sequencing Facility for performing the Illumina RNA-seq. We thank Christina Ochsenbauer for providing the subtype C HIV-1 molecular clones. This study was supported by the Intramural Research Program of the Center for Cancer Research, National Cancer Institute, NIH, and by an intramural AIDS Research Fellowship (for M.V.C.). J.F.H. is supported by the Gilead Sciences Research Scholars Program in HIV, NIH/NIGMS funding for the HIV Accessory & Regulatory Complexes (HARC) Center (P50 GM082250), NIH funding for the Third Coast Center for AIDS Research (P30 AI117943), National Health and Medical Research Council Idea Grant 2013215, and NIH/NIAID grants for HIV research (R01 AI150455, R01 AI165236, and R01 AI150998). S.B.V.E. is supported by NIH/NIAID grant for HIV assembly research (R01 AI138625)

## Author Contributions

M.V.C. and E.O.F. designed the study, interpreted the data, and wrote the manuscript. M.V.C., H.K.H., H.C., and L.M.S. performed the experiments. H.K.H. designed the guide RNA targeting FIP1C and generated the lentivector used to generate the SupT1huR5 and H9 FIP1C KO cell lines. H.K.H. also generated the Jurkat E6.1 FIP1C KO clonal cell lines and the H9 FIP1C KO cells. M.V.C. generated the SupT1huR5 FIP1C knockout cell line, performed all Env incorporation and spreading infection assays. H.C. and L.N. assisted with spreading infection assays. L.M.S. and J.F.H. performed the virus replication assays in the FIP1C KO primary CD4^+^ T cells. D.A.S., J.F.H., S.B.V.E. contributed valuable feedback and guidance on CRISPR/Cas9 KO and trafficking experiments. S.A. isolated hPBMCs from human blood and stimulated them for spreading infection assays. L.N. assisted with cell density comparisons in spreading infection assays. All authors contributed to preparation and editing of the manuscript.

